# Individual differences reveal shared computational structure in perceptual decision-making

**DOI:** 10.64898/2026.11.04.764343

**Authors:** Jingming Xue, Samuel F. Feng, Hua-Dong Xiong, Shensheng Wang, Robert C. Wilson

## Abstract

To discover computational mechanisms of cognition, models typically specify a fixed psychological architecture, treating individual differences as variations within a predefined parameterization. Here, we take a different approach by using the population structure of individual differences to directly drive model discovery. We develop a low-rank neural network that uses this ‘population geometry’ to learn shared computational components and participant-specific weights from trial-level choices, allowing cross-subject variation to constrain the model’s architecture. Applied to perceptual decision-making in the Bernoulli Clicks Task across primary and replication datasets, out-of-sample prediction reveals a six-dimensional representation for individual differences: two continuous dimensions of temporal evidence weighting and two of choice-outcome history dependence, alongside participant-specific side bias and signal-to-noise ratio. This low-dimensional organization replicated in an independent dataset. Together, these findings reveal that seemingly disparate decision heuristics reflect different weightings over a small set of shared computational components, and establish a principled methodology for using behavioral variability to discover the mechanistic architecture of cognition.

## 1 Introduction

The study of general cognitive mechanisms and the study of individual differences have largely developed as separate traditions. Research aimed at explaining cognition typically prioritize canonical, population-averaged computations—often treating individual variability as measurement noise to be averaged out (or parameter variations across subjects). In contrast, differential psychology and related traditions place individual differences at the center of inquiry, documenting systematic variation in cognitive abilities, behavioral tendencies, and psychological traits. However, these differences are often characterized through descriptive statistical dimensions without specifying the computational mechanisms that generate them. Consequently, general accounts of cognition often leave individual differences unexplained, while accounts of individual differences frequently lack a mechanistic foundation.

This divide persists even within computational cognitive modeling. Traditional cognitive models specify a set of psychologically interpretable computations, and fit their interpretable parameters, such as evidence sensitivity or learning rate, to each participant’s trial-by-trial behavior [17, 18, 32]. Individual differences must therefore be expressed within that predefined parameterization. Neural-network models provide greater flexibility by allowing computations to emerge through learning [2], but individual differences are still commonly examined after training, through post hoc analyses of model representations or participant-specific parameters. Although these approaches can characterize how individuals differ within a model, the structure of individual differences rarely plays a direct role in determining which computational mechanisms the model should learn. Individual variability is therefore still examined primarily after a model has been constructed, rather than as a source of information that can guide the discovery of the model itself.

We propose that the geometry of individual differences can directly constrain the discovery of cognitive mechanisms. Here, population geometry refers to how participant-specific low dimensional representation are organized relative to one another. That is, variation across people comes from combining a few shared computation components in different proportions. Because each person’s behavior is generated by weighting these same components differently, systematic relationships across individuals should reflect their common computational structure. Individual differences therefore do not merely represent parameter variations to be characterized after a model is constructed; they instead reveal the underlying dimensions that characterize a cognitive process.

The ABO blood-group system provides a concrete example of how population-level patterns of individual differences can reflect underlying biological mechanisms. When red blood cells from one individual are mixed with serum from another, some combinations produce agglutination while others do not. Importantly, these reactions are not independent or arbitrary: they exhibit systematic relationships across individuals. For example, serum containing anti-A antibodies agglutinates red blood cells from both type A and type AB individuals because both carry the A antigen. These patterns, which emerge at the macroscopic level in the correlational structure across blood samples, can be explained by a small set of microscopic components: the presence or absence of A and B antigens on red blood cells and their corresponding antibodies in serum [8]. The same biological components thus explain both individual reactions and the structure of variation across the population.

We seek an analogous connection in cognitive modeling: identifying computations that explain both choices on individual trials and systematic relationships among behavioral features across people. We implement this approach using a neural network in which individual decision-making is expressed through a small set of shared computational components. Participant-specific coefficients, hereafter termed participant scores, determine how strongly each component contributes to an individual’s choices. The model learns the shared components and participant-specific coefficients jointly from trial-by-trial behavior across the population. Therefore, the model allows the structure of individual differences to constrain the discovery of shared cognitive computations by requiring individual decision-making to be expressed through a common low-dimensional basis.

We apply this approach to perceptual decision-making, where optimal behavior can be specified explicitly, yet individuals differ systematically in how they use information to make choices [5, 17, 28]. In the Bernoulli Clicks Task, participants report which ear received more clicks [17, 18]. Optimal behavior requires counting clicks equally regardless of their temporal position, without relying on previous choices or outcomes. Human decisions depart from this rule in diverse ways: participants place different weights on early and late evidence [6, 23, 38, 40], and differ in how previous choices and outcomes influence their current decisions [1, 34]. These diverse patterns of suboptimality may not require equally diverse computational components. Instead, they may arise from different participant-specific combinations of a smaller set of shared components. We investigate this possibility by using the structure of individual differences to constrain the dimensionality of individual variation in temporal weighting and history dependence using trial-by-trial choices.

Temporal weighting provides a first example of how diverse suboptimal behaviors may reflect a shared computational structure. Previous studies have shown that different temporal weighting kernels can arise from common computational mechanisms. For example, dynamic divisive normalization can generate different weighting kernels through changes in its parameters [17], while hierarchical approximate inference provides an account of both primacy and recency effects [22]. However, demonstrating that a model can generate different weighting kernels does not establish how these kernels are related across individuals. In particular, it remains unclear whether the observed variation in temporal weighting can be captured by a small number of shared dimensions. We therefore examine whether individual weighting kernels can be explained by a low-dimensional representation learned directly from trial-by-trial choices, rather than treating different kernels as independent behavioral strategies.

History dependence provides a complementary example. Previous studies have identified several systematic influences of past decisions on current choices, including outcome-dependent win–stay/lose–shift tendencies and outcome-independent choice repetition or alternation [1, 18, 21, 35]. These effects have often been associated with distinct computational processes, such as reinforcement learning [11] and choice-history biases. However, distinguishing these effects at the behavioral level does not necessarily imply that they represent independent dimensions of individual variation. Their strengths may covary systematically across participants, reflecting a common underlying component, or vary independently, requiring additional dimensions. We therefore examine whether the structure of individual differences supports the conventional separation of history-dependent influences or reveals a different low-dimensional organization.

To investigate these questions, we systematically vary the number of shared components used to represent temporal weighting and history dependence, evaluating each model’s ability to predict held-out choices. We then examine the learned components and participant-specific coefficients to characterize the dimensions of individual variation and their relationships to observed behavioral patterns. Finally, we assess whether the identified structure is reproduced in an independent replication dataset. Together, these analyses test whether the diversity of suboptimal decision-making can be captured by a small number of shared dimensions that account for both trial-level choices and systematic variation across individuals.

## 2 Individual Differences in a Perceptual Decision Making Model

Participants performed the Bernoulli Clicks Task,^17,18^ as illustrated in Fig. 1. Each trial began with a 1-s fixation period, followed by 20 auditory clicks presented over 1 s, with one click delivered to either the `Left` or `Right` ear every 50 ms. On each trial, a target side was randomly assigned with equal probability (50%). Each click was independently delivered to the target ear with probability *p* = 0.55 and to the opposite ear with probability 1*− p* = 0.45. Participants indicated which ear they judged to have received more clicks by pressing the corresponding `Left` or `Right` key. Following each response, an arrow indicating the selected side was displayed for 500 ms, followed by accuracy feedback for 500 ms (1 = correct, 0 = incorrect).

**Figure 1.**
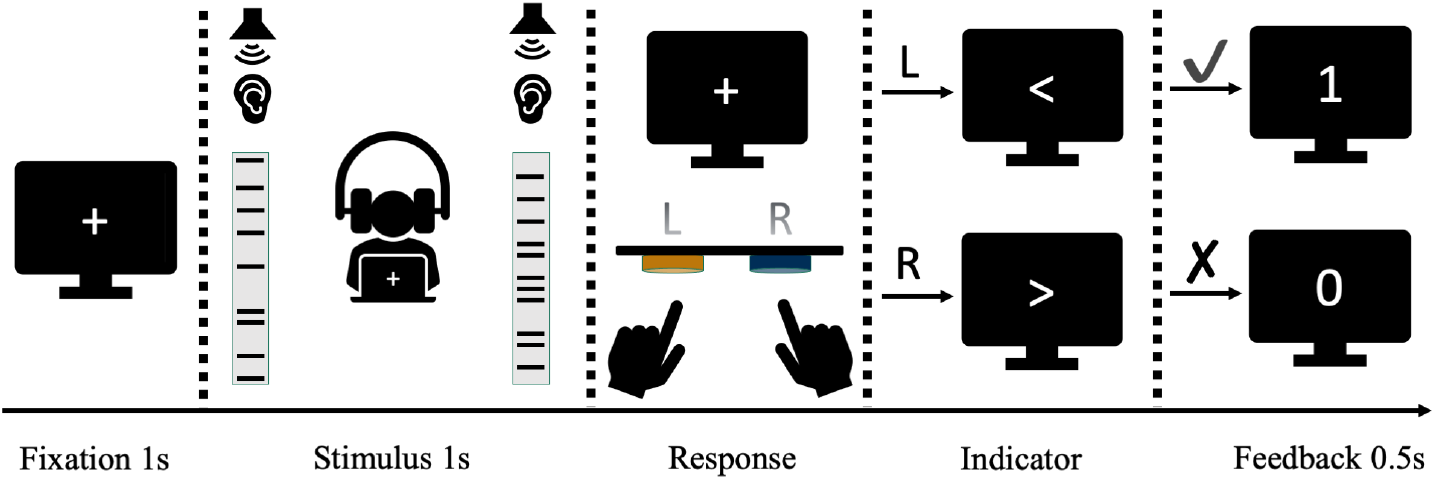
Task procedure. Participants listened for 1 s to a sequence of 20 auditory clicks, each presented to either the `Left` or the `Right` ear, and then reported which side had received more clicks. The 1-s stimulus interval was divided into twenty 50-ms bins, with exactly one click occurring in each bin on either the `Left` or the `Right` side.

### 2.1 Optimal model

The optimal thing to do in the Bernoulli Clicks Task is straightforward - simply count the clicks on each side and choose the side with most clicks.^17^

Written mathematically, the optimal model has a particularly simple form that serves as a baseline model to which suboptimalities can be added to account for human behavior.

Specifically, if we write the sequence of twenty clicks on trial *t* as,

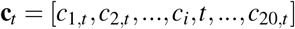

and define the *i*th click in the click train to be,

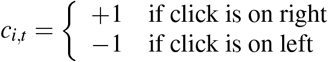

then the optimal model simply sums these terms into a decision variable, *DV*_*t*_, and determines its sign (+, *−*). In notation, the optimal model first computes:

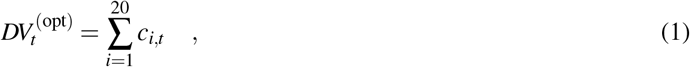

and then determines the choice probability from the sign of 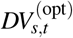, breaking the tie with a 50-50 coin flip when 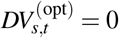. Letting α_*t*_ *∈ {−*1, +1*}* denote the choice on trial *t*, the optimal behavior thus results in:

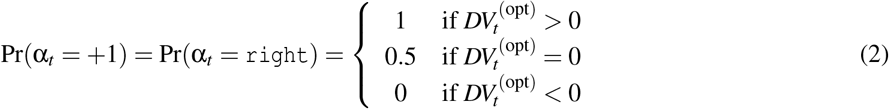

With a view towards extending this to human behavior, we can rewrite this choice rule using a softmax function:

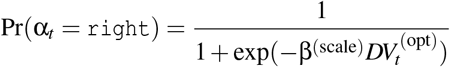

where β^(scale)^ is an inverse temperature parameter. Equation 2 is recovered when β^(scale)^ = +∞.

The behavior of the optimal model is characterized by its psychometric choice curve - the probability of choosing `Right` as a function of the difference in clicks between `Left` and `right`. As shown in Fig. 2A, this leads to a step function choice curve consistent with the deterministic behavior of the optimal model when 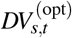 is non-zero.

**Figure 2.**
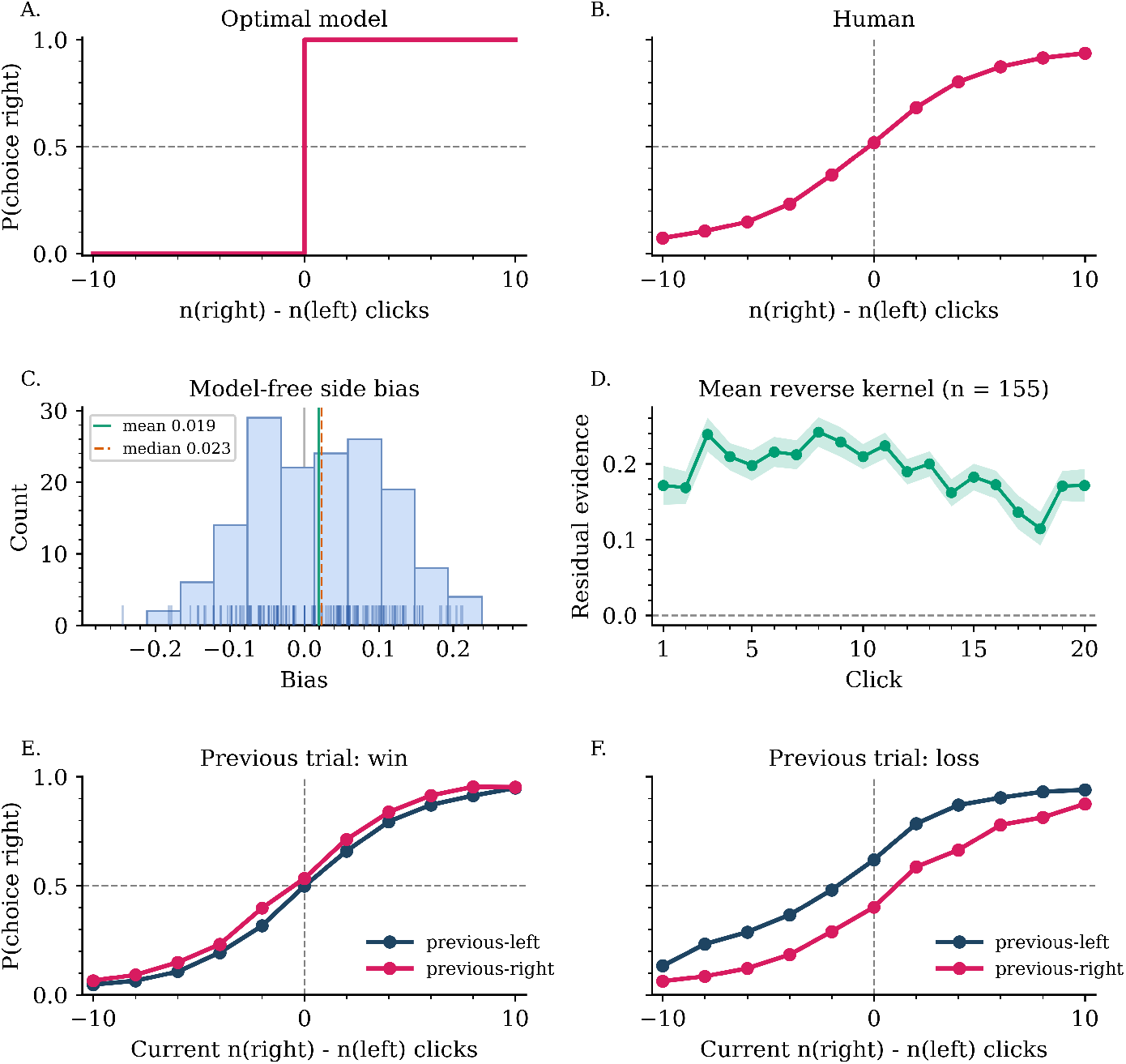
Suboptimal behavior deviations from the optimal model in the Bernoulli Clicks Task. **A**, The optimal model chooses the `Right` when there are more clicks on that side, producing a step-like psychometric function. **B**, Instead, human subjects from the first dataset shows a sigmoid shaped psychometric function, the probability vary more gradually with accumulated evidence. **C**, Distribution of the side bias from human subject, showing substantial variability. **D**, Example psychophysical reverse kernel over all participant. At each click position, the kernel illustrates how much an unexpected sensory-evidence fluctuation at each moment predicts the eventual decision. It therefore characterizes the behavioral sensitivity to unexpected evidence fluctuations beyond the stimulus baseline. The shaded area is the pointwise 95% confidence interval for the mean reverse kernel across the 155 subjects. **E**, Choice curves conditioned on the previous trial being win/correct, showing a tendency to repeat the previous choice after a win. **F**, Choice curves conditioned on the previous trial being loss/incorrect, showing a tendency to switch away from the previous choice after a loss.

### 2.2 Humans show different suboptimalities on the clicks task

Human performance is suboptimal on the Bernoulli Clicks Task. This is easily seen by plotting the choice curve for people in Fig. 2B. In contrast to the optimal model, people make mistakes, sometimes choosing `Left` when there were more clicks on the `Left` and choosing `Right` when there were more clicks on the `Right`.

The simplest way to model these mistakes is to assume that, instead of computing the correct decision variable, people compute a noisy version.^9^ This noise has the effect of changing the inverse temperature parameter on the optimal model from +∞ to a finite number. That is, for human subject *s*, the corresponding inverse temperature parameter satisfies 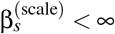. This gives an expression for the choice probabilities of subject *s* as

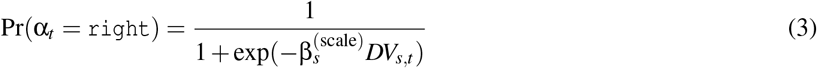

While this simple model of suboptimal human behavior provides a good fit for the overall choice curves, it assumes that all human mistakes are completely random. This is not the case [3]. People show a preference for one side or the other when there was equal evidence for both sides [20] (Fig. 2C).

Averaged across people, this side bias is close to zero, but individual participants can exhibit a preference for either `Left` or `Right` choices (Fig. 2C). Such a bias can easily be accommodated in the model by adding an intercept term, 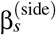 to the decision variable:

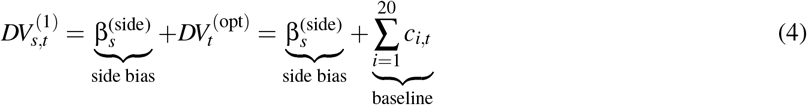

with 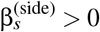 corresponding to a `Right` bias and 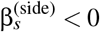 corresponding to a `Left` bias.

Second, people do not give each click the same weight in their decision.^17^ A simple way to illustrate this is with the choice-triggered reverse kernel (Fig. 2D),which measures, at each click position, how unexpected fluctuations in sensory evidence relative to the trial’s expected stimulus statistics are associated with the eventual choice. The fluctuations of this kernel indicate uneven behavioral sensitivity to evidence over time. This uneven weighting of clicks can be accommodated by adding click-specific deviations from the optimal weights to the decision variable,

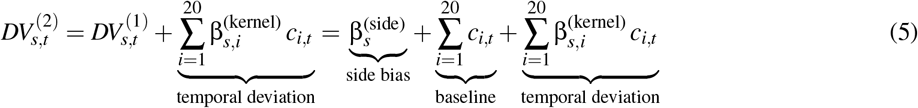

Here, 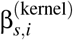 captures the extent to which the weight assigned to click *i* differs from its optimal value, with 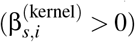 indicating overweighting and 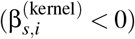 indicating underweighting relative to the optimal model.

Finally, people are influenced by their history of past choices and outcomes.^1,18^ These order effects are easily seen by computing the choice curves conditioned on the last choice and outcome (Fig. 2E & F). In the Bernoulli Clicks Task, these history effects correspond to a small win-stay effect, in which people tend to repeat their last choice when it was correct, and a larger lose-shift effect, in which people tend to choose the alternate option when their last choice was wrong. Expanding the model to include such history effects leads to an updated decision variable of the form

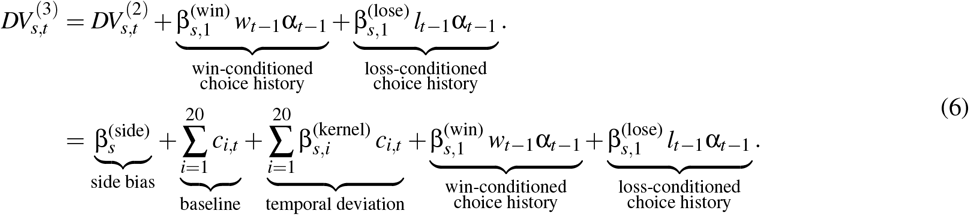

where α_*t−*1_ *∈ {−*1, +1*}* denotes the choice on the preceding trial, coded as +1 for a rightward choice and *−*1 for a leftward choice. and *w*_*t−*1_ and *l*_*t−*1_ are indicator variables that code whether the last choice was a win (*w*_*t−*1_ = 1, *l*_*t−*1_ = 0) or a loss (*w*_*t−*1_ = 0, *l*_*t−*1_ = 1). 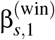 and 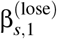 describe the effects of win-conditioned and loss-conditioned choice history from the last trial for subject *s*. Stay or shift tendencies are determined by the signs of these fitted coefficients.

Of course, history effects may persist beyond just the last trial. To account for this possibility we can extend the history kernel to include 5 past trials.

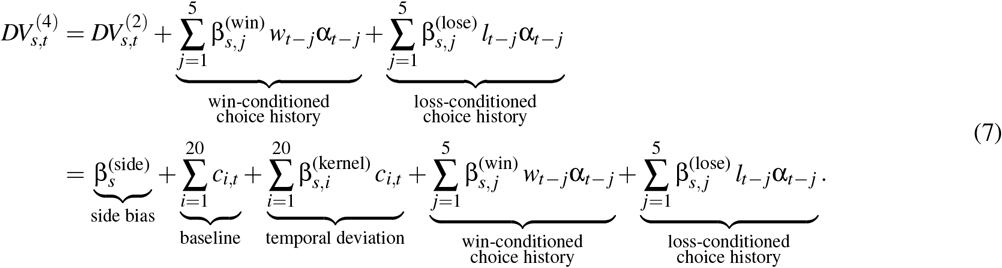

where 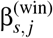 and 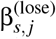 define the history kernels for subject *s*, capturing how choices and outcomes from *j* trials in the past influence the current decision.

### 2.3 Average Model Behavior Across participants

Our model closely resembles the logistic model of human performance on the Bernoulli Clicks Task used in Keung et al.^18^ Fitting this model to behavior allows us to characterize the behavior of each individual subject, *s*, via 32 free parameters: 1 evidence-scaling parameter, 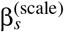, which captures the overall sensitivity to accumulated evidence; 1 side-bias parameter,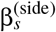; 20 temporal kernel deviation parameters, 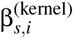, which describe how the weight assigned to click *i* deviates from uniform evidence accumulation; and 10 history parameters, comprising five win-history weights, 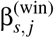, and five lose-history weights, 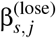, for *j* = 1, …, 5. Together, these parameters quantify the extent to which each subject’s behavior departs from the optimal observer in terms of overall evidence sensitivity, uneven click weighting, sequential dependencies, and side bias.

Taking the average of these parameters across 155 participants (see §5.1), we obtain the results shown in first row of Fig. 3. The mean scaling parameter, 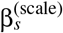, was below 1, indicating that the decision variable was attenuated before being mapped to choice probability. A *Greenhouse–Geisser corrected repeated-measures ANOVA* provided evidence for an uneven temporal deviation: integration weights differed significantly across the 20 click positions,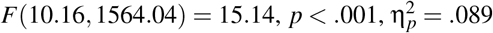. We also observed a reliable right-side bias: a *one-sample Wilcoxon signed-rank test* showed that side-bias coefficients were significantly greater than zero, *W* = 8410.00, *p <* .001, rank-biserial *r* = .391, *M* = 0.409, *Mdn* = 0.312. Finally, sequential effects were evident over several trials into the past. Averaging coefficients across lags 1–5, a *one-sample Wilcoxon signed-rank test* indicated that the overall win-stay effect was significantly greater than zero, *W* = 10620.00, FDR-corrected *p <* .001, rank-biserial *r* = .757, *M* = 0.270, *Mdn* = 0.211, whereas a second *one-sample Wilcoxon signed-rank test* showed that the overall lose-shift effect was significantly less than zero, *W* = 2481.00, FDR-corrected *p <* .001, rank-biserial *r* = *−*.590, *M* = *−* 0.353, *Mdn* = *−* 0.197. Interestingly, after the first two trials into the past, the lose-related coefficients became positive, suggesting that more distant loss history may reflect a more general stay effect rather than a local lose-shift tendency.

**Figure 3.**
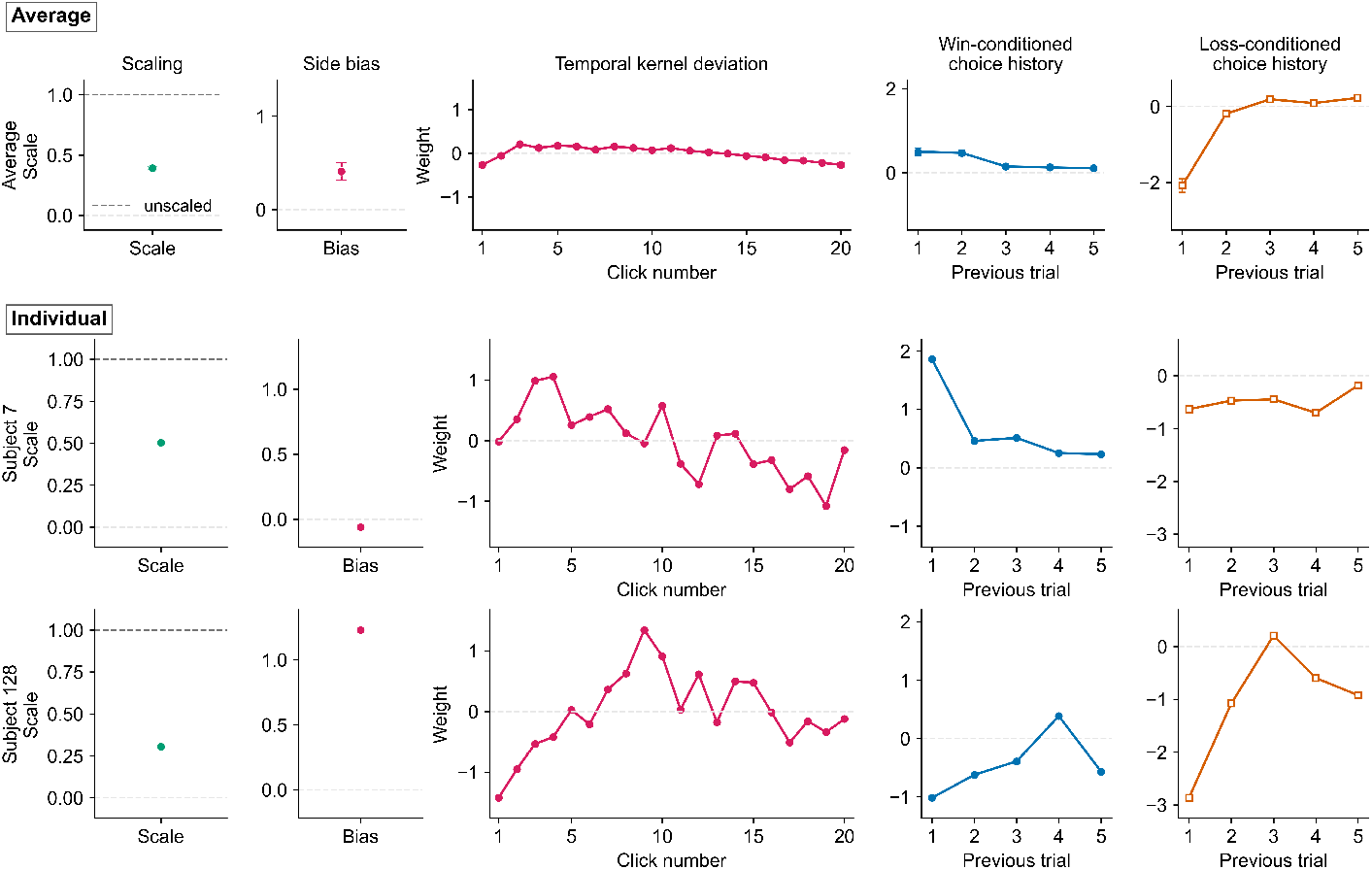
Regression derived average and individual differences in suboptimalities. Each row summarizes the fitted parameters for one level of analysis: the top row shows the mean coefficient across participants, and the remaining rows show two representative subjects. The first column shows the scaling parameter; the dashed line horizonally at 1 denotes no scaling. The second column shows the subject-specific side-bias coefficient, where positive values indicate a rightward bias and negative values indicate a leftward bias. The third column shows the temporal deviation, that is, the deviation in the weight assigned to each of the 20 clicks from the baseline. The fourth and fifth columns show the win-history and loss-history kernels, respectively, across the previous five trials. Positive win-history coefficients denote repeating a previously rewarded choice (win–stay), whereas negative loss-history coefficients represent switching after an error (lose–shift). Together, these panels illustrate the heterogeneity across individuals in temporal evidence weighting, side bias, sequential dependencies, and overall decision sensitivity.

### 2.4 Individual differences in suboptimalities

Example subject-level kernels reveal substantial individual variability in the structure of choice suboptimalities. The first column in Fig. 3 shows the weight of the recovered temporal deviation for the selected 2 subjects. The temporal deviations show that subjects differ in how they weight clicks across time, one subjects over-weighting early (e.g., Subject 7) clicks, another subject show largest relative weight in the middle (e.g., Subject 128).

The history kernels show equally marked individual variability in sequential choice biases, a positive win coefficient indicates win-stay behavior, whereas a negative win coefficient indicates win-switch behavior. For losses, the interpretation is reversed relative to the common verbal label: a negative loss coefficient corresponds to lose-shift behavior, while a positive coefficient corresponds to lose-stay behavior (See the third and fouth columns of Fig. 3). We classified the subjects into 4 category based on the sign of the mean weight from the past trials. For example, Subject 7 shows standard negative win-stay coefficient with all win coefficients being negative, and positive loss effects with loss coefficients all negative. Subject 128 is classified as alternation. It shows strongly negative first-lag win and loss coefficients, indicating pronounced immediate shifting after both wins and losses. Although one win coefficient and one loss coefficient become positive at intermediate lags, the overall pattern remains negative, consistent with a broader alternation tendency.

The side-bias and scaling terms further show that these individual differences are not limited to history effects and temporal deviations. As shown in second column of Fig. 3. Subject 128 has a strong positive coefficient, indicating a baseline tendency to the `Right` side; Subject 68 has a negative coefficient, indicating a baseline tendency to the `Left` side. Similarly, the scaling parameter also varies across subjects, reflecting differences in overall sensitivity to the decision variable. For example, Subjects 75 show larger scaling values than others, indicating stronger overall weighting of the model’s decision variable. Importantly, the effective contribution of each suboptimality depends on both its raw coefficient and the subject-specific scaling parameter. Thus, a large raw coefficient can be attenuated by a small scaling value. For example, Subject 128 combines a lower scaling value with a strong positive side bias, yielding an effective side-bias magnitude that is more comparable to Subject 68 despite their different raw coefficients.

Together, these examples show that suboptimalities vary along multiple axes, including integration-kernel deviations, side bias, and history effects. This motivates the central question of this current research: whether these subject-specific suboptimalities can be represented in a shared low-dimensional space across individuals.

### 2.5 Constraining mechanism discovery with a low-rank neural network

We first fit the noisy optimal model with net click evidence and participant-specific scaling (Fig. 4, blue), then add participant-specific side bias (green). These models provide baselines for introducing low-rank factorizations of the temporal and joint history coefficient matrices. In each factorization, shared components and participant-specific weights are learned jointly from trial-level choices across the population, allowing individual differences to constrain the learned computations.

**Figure 4.**
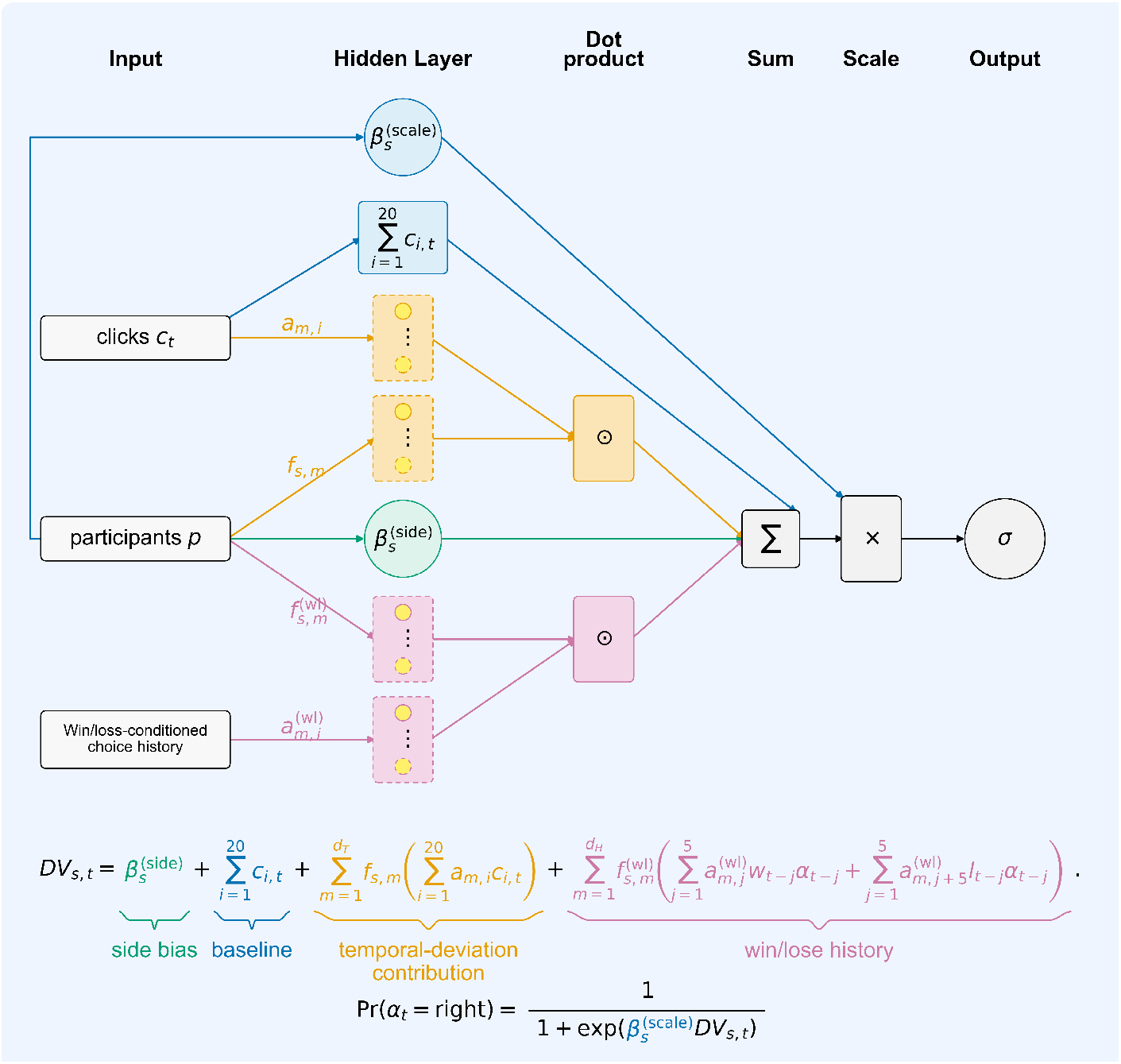
Low-rank neural-network model for individual differences in the Bernoulli Clicks Task. On each trial, the model combines three sources of information: the 20-click evidence sequence *C*_*t*_, the participant identity *p*, and recent win/loss history. The click sequence contributes a baseline accumulated-evidence term, 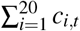, and is also projected onto shared temporal components *a*_*m,i*_. Participant-specific scores *f*_*s,m*_ weight these basis projections, producing an individualized temporal deviation. In parallel, participant identity provides a side-bias term 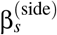 and a positive evidence-scaling parameter 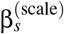. Win- and loss-conditioned choice histories are projected onto shared history components 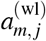, whose contributions are weighted by participant-specific history scores 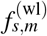. The temporal-integration term, history term, baseline evidence, and side bias are summed to form the decision variable *DV*_*s,t*_, which is then multiplied by the subject-specific scaling parameter and passed through a logistic function to predict the probability of a rightward choice. This architecture represents each participant’s temporal kernel deviations and joint history kernels as linear combinations of shared components, preserving individual variability while constraining it to a low-dimensional space.

One approach to identifying shared components in temporal weighting is to estimate each participant’s temporal deviation using Eq. (7) and then apply factor analysis to the resulting coefficient matrix. However, this two-stage approach estimates individual kernels independently of their shared structure. We instead learn the shared temporal components and participant-specific weights directly from trial-level choices across the population by imposing the following factorization:

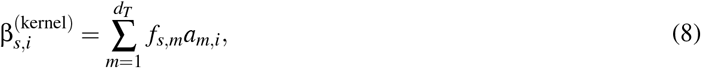

Here, *d*_*T*_ is the temporal latent dimensionality. Each dimension *m* is represented by a shared temporal component **a**_*m*,:_, whose entry *a*_*m,i*_ is the weight at click position *i*. The participant score *f*_*s,m*_ gives subject *s*’s coordinate on that dimension. The components serve as basis functions for reconstructing participant-specific temporal kernel deviations. Summarizing over 20 weight and coupling with the clicks information, we factorize the temporal deviation in Eq. (5) into:

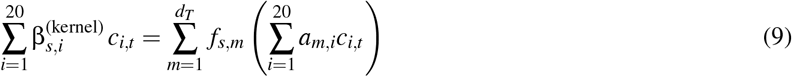

So the Eq. (5) can be rewritten as:

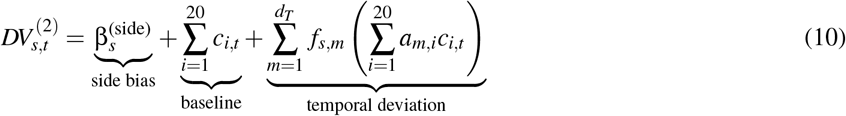

This model is equivalent to a feedforward neural network with the following form, blue, yellow and green part in Fig. 4.

We then trained the model to select the temporal latent dimensionality *d*_*T*_ using validation loss. The training process is a method similar to the stacked RBMs or greedy layer-wise pretraining,^4^ but differs in that we repeat the training single-neuron-wise rather than layer-wise (See Method Algorithms 1 for training details). The train/validation splits are fixed across *d*_*T*_, and the loss *ℒ* is the binary cross-entropy evaluated on the validation set (See Section 5.5 for more training details).

As with the temporal coefficient matrix, we represent the joint history coefficient matrix using a low-rank factorization. For the win-conditioned choice-history term, we write

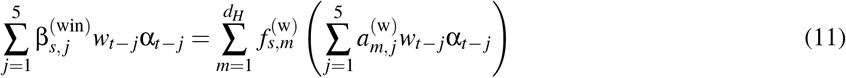

Similarly, for the loss-conditioned choice-history term, we write

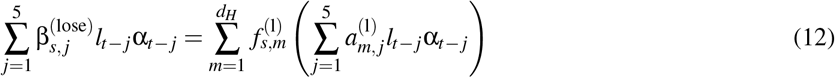

We use shared participant-specific weights to combine the win- and loss-conditioned choice-history terms:

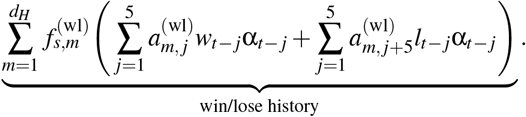

Here, the superscript (wl) denotes the combined win–loss history kernel. The history dimensionality is *d*_*H*_. The index *m* = 1, …, *d*_*H*_ indexes a latent history dimension that is shared by the win-conditioned and loss-conditioned choice-history terms, and 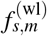 is subject *s*’s score on this joint history dimension.

For each latent dimension *m*, the first five coefficients, 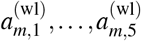, specify the weights on win-conditioned choice history over lags *j* = 1, …, 5. The next five coefficients, 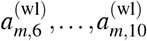, specify the corresponding weights on loss-conditioned choice history. Thus, 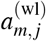 and 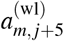 refer to the win and loss portions, respectively, of the shared history component corresponding to dimension *m*,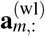. In addition, the single score 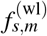 scales both the win and loss portions of latent component *m* together. Therefore, the model represents each subject’s history dependence as a combination of *d*_*H*_ joint win–loss components, rather than as two independently scaled win and loss kernels.

The final form of the model to select the latent dimensionality of history dependence is the full model in (Fig. 4):

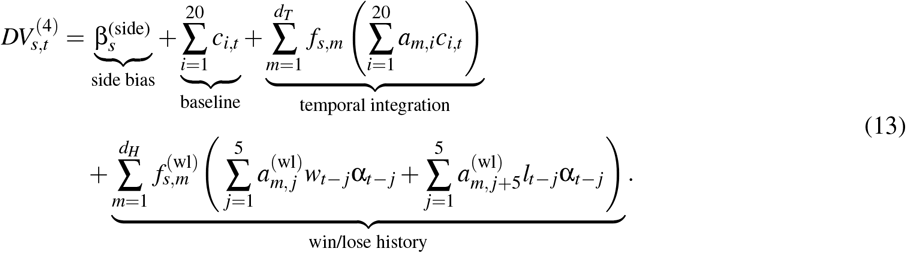

Similarly to the temporal deviation training, we trained the model to select the latent dimensionality *d*_*H*_ of history dependence using validation loss. (See Method Algorithms 2 for training details). The train/validation splits are still fixed across *d*_*H*_, and the loss *ℒ* is the binary cross-entropy evaluated on the validation set (See Section 5.5 for more training details).

## 3 Results

### 3.1 Side biases and Decision noise

We obtain the results from the final model (i.e., Eq. (13)). Individual participants show varying `Left`–`Right` biases (see `Left` panel in Fig. 6). A two-sided Wilcoxon signed-rank test indicated that subject-level bias differed significantly from zero, *W* = 3653.00, *p <* .001, *Mdn* = 0.193, *IQR* = 1.182, *N* = 155. At the individual level, 96 participants exhibited a positive (`Right` side) bias estimate and 59 exhibited a negative (`Left` side) bias estimate. For scaling factor, all estimated scaling factors satisfy 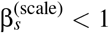, with a median of 0.354 (IQR = 0.203; *Q* = 0.255, *Q*_3_ = 0.459), indicating generally attenuated stimulus–choice sensitivity but between-subject variability (See `Right` panel in Fig. 6).

### 3.2 temporal deviation exhibits all types of kernel shapes across individuals

Validation loss favored a two-dimensional representation of individual differences in temporal weighting under the Bernoulli Clicks Task. As shown in the `Left` panel of Fig. 5, the average validation loss reaches its minimum when the model has *d*_*T*_ = 2 neurons in both the click and subject layers. By constraining the latent space to *d*_*T*_ = 2, the model significantly reduces the required parameters for the temporal kernel from 3100 to 346, thereby acting as a structural regularizer that prevents overfitting while still effectively capturing trait variation (see Section **??**).

**Figure 5.**
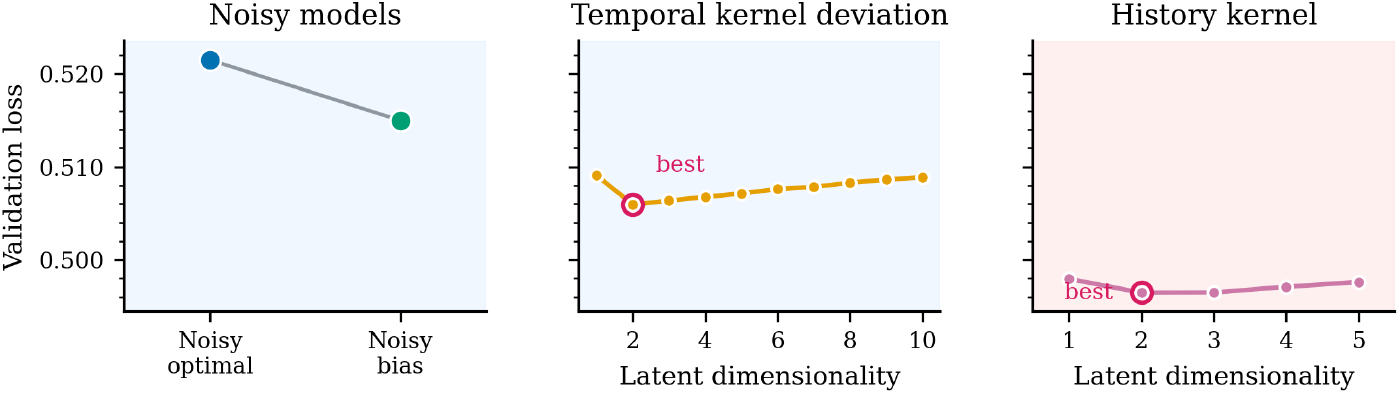
Validation loss for all models. `Left` panel show validation loss for Noisy optimal model and Noisy bias model. Middle and `Right` panel show validation loss as a function of the number of latent dimensions for the temporal weighting kernel (Middle) and the history kernel (Right). Each point shows the average loss on the validation set over 121 training loops. In both cases, the minimum loss occurs at two latent dimensions, indicating a best-fitting latent dimensionality of 2 for both temporal weighting and history dependence.

**Figure 6.**
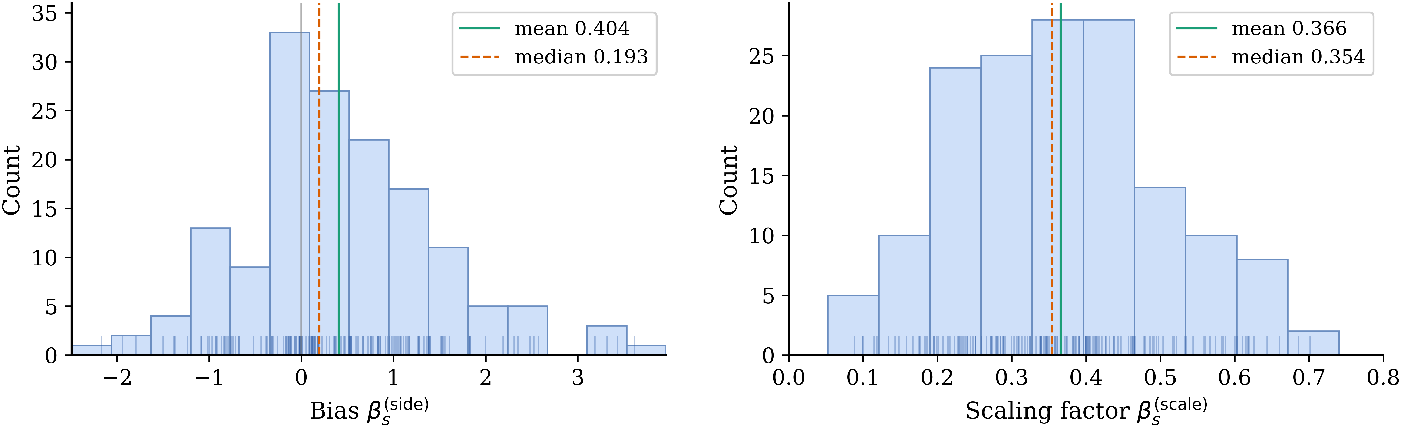
Distribution of estimated choice bias across participants. Bars show the histogram of individual bias parameters, with rug marks indicating single participants. The solid vertical line denotes the mean bias (0.404); the dashed line denotes the median bias (0.193).

Individual differences in temporal weighting are captured by two latent dimensions, each represented by a shared temporal component (Fig. 7a). Participant scores give coordinates in this two-dimensional space (Fig. 7b). The first dimension separates participants who emphasize late evidence from those who overweight mid-trial evidence. High positive scores on this dimension correspond to a bump-shaped kernel that emphasizes information near the middle of the trial (Fig. 7a, blue). In contrast, strong negative scores correspond to a recency effect, in which decisions are driven primarily by the latest evidence. The second dimension captures a primacy effect: high positive scores correspond to kernels that strongly overweight the earliest evidence (Fig. 7a, yellow). Thus, each participant’s temporal kernel deviation is reconstructed by combining the components corresponding to the two dimensions:

**Figure 7.**
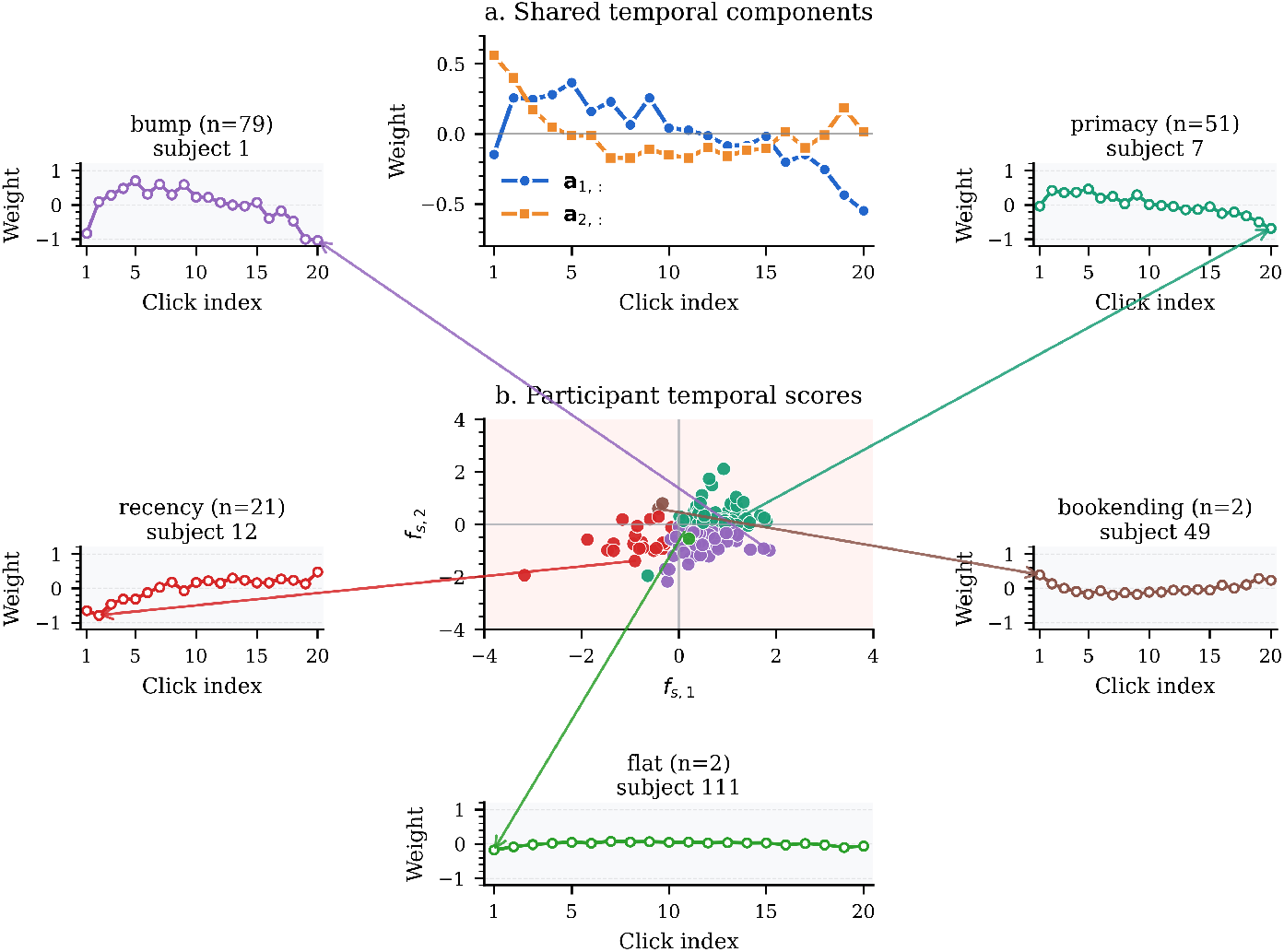
Shared temporal components, participant scores, and individual temporal kernel deviations in a two-dimensional representation. (a) Per-click weights for the two shared temporal components learned by the model. (b) Each point corresponds to a participant, plotted by their scores *f*_*s*,1_ and *f*_*s*,2_ on the two temporal-kernel dimensions. Colors indicate clusters associated with distinct temporal weighting kernels. (b, callouts) Example temporal kernel deviations from individual participants representative of each cluster: bump-shaped, primacy, recency, bookending, and flat. Callout titles report the number of participants assigned to each pattern.

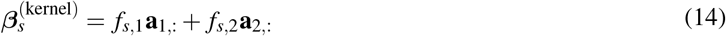

where **a**_1,:_ is a vector with entries **a**_1,*i*_ over 20 clicks, representing the shared temporal component corresponding to the first latent dimension (Fig. 7a, blue). *f*_*s*,1_ is the corresponding score at this dimension for individual *s*; and **a**_2,:_ is the shared temporal component corresponding to the second latent dimension (Fig. 7a, blue), *f*_*s*,2_ is the second dimension’s score for individual *s*.

This low-rank framework reconstructs the full spectrum of observed weighting strategies. Subjects indicated by the blue points are mostly clustered in the lower-`Right` quadrant and show a bump shape: evidence in the middle of the sequence is weighted most strongly, with reduced weighting of the earliest and latest clicks (Fig. 7c). When both *f*_*s*,1_ and *f*_*s*,2_ scores are positive, the resulting kernels overweight early clicks and gradually downweight later ones, corresponding to a primacy or “early decider” strategy (Fig. 7d). Strongly negative *f*_*s*,1_ scores produce the opposite pattern: early evidence is underweighted and late evidence is emphasized, yielding a recency or “procrastinator” strategy (Fig. 7e). Furthermore, two participants exhibited a “bookending” strategy, characterized by negative *f*_*s*,1_ scores and positive *f*_*s*,2_ scores. These participants weighted the beginning and end of the evidence stream strongly, while downweighting evidence presented in the middle. Finally, subjects whose *f*_*s*,1_ and *f*_*s*,2_ scores both lie near zero produce kernels with low variance and nearly zero slope, corresponding to an approximately flat, near-optimal weighting pattern (Fig. 7g).

### 3.3 Two dimensions characterize individual differences in history dependence

Model comparison indicated that the optimal latent dimensionality of the history kernel was *d*_*H*_ = 2.(Fig. 5, `Right` panel). In addition, with one dimension denoting the side bias, and one dimension denoting the signal-to-noise ratio, the individual variation under the current data structure can be explained by six dimensions.

Individual differences in history kernels are captured by two latent dimensions (Fig. 8a). The component corresponding to the first dimension, 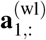, embodies a classic Reinforcement Learning (RL) strategy. It assigns opposite weights to the outcomes of the previous trial: a positive weight following a win and a negative weight following a loss. A positive score on this dimension thus reflects a strong “Win-Stay, Lose-Shift” tendency. The component corresponding to the second dimension, 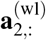, captures an outcome-independent Choice Bias. It assigns a strong weight to the previous action regardless of its outcome, reflecting an intrinsic preference to either repeat a choice (perseveration) or switch between choices (alternation). Each participant’s joint history kernel is reconstructed as:

**Figure 8.**
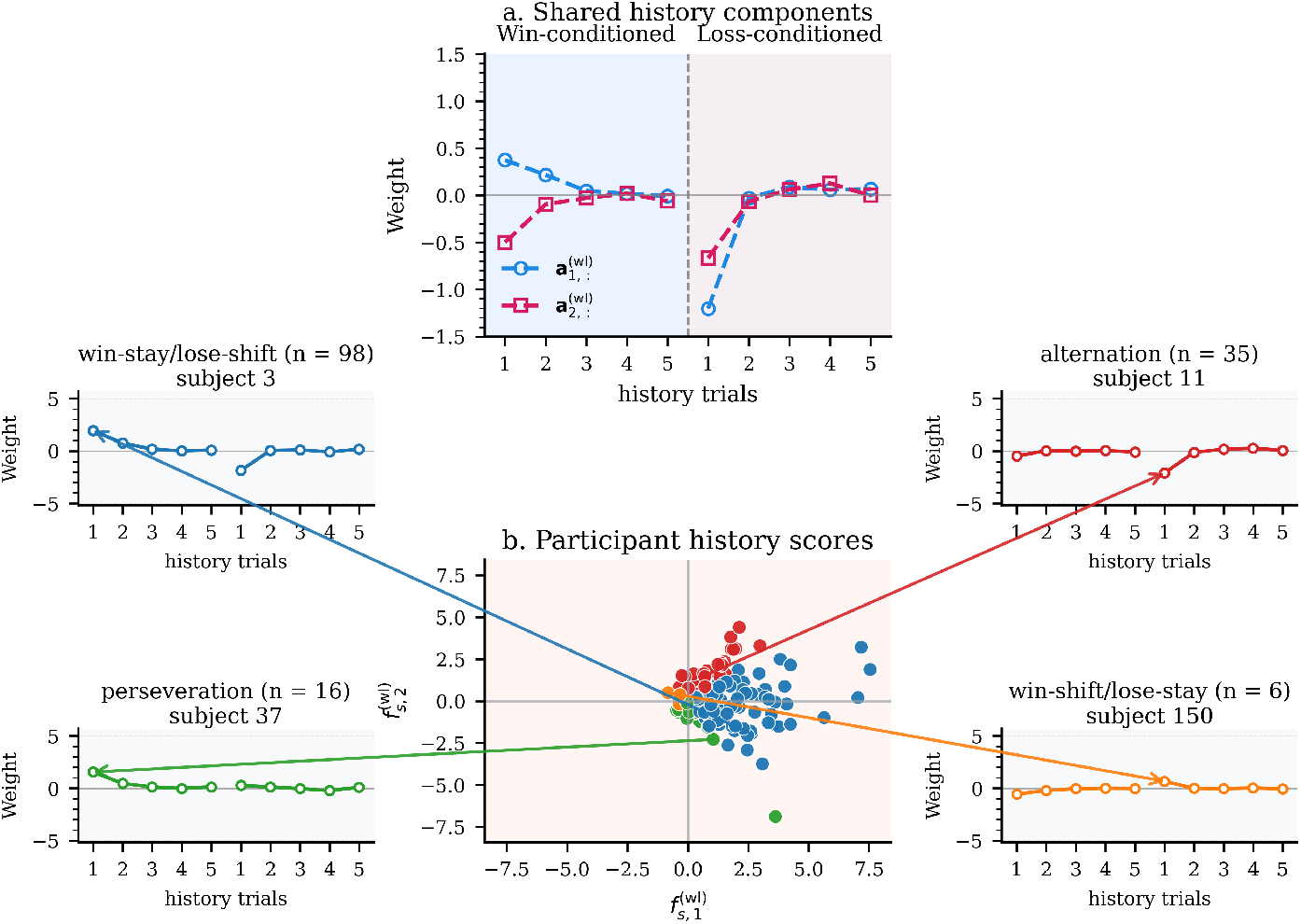
Two-dimensional history space and example history kernels. (a) Weights over the preceding five trials for the win-conditioned (`Left` panel) and loss-conditioned (`Right` panel) portions of the shared history components. (b) Each point is a participant plotted by their 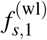 and 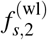 scores; colors indicate clusters corresponding to distinct history strategies. (b, callouts) Example kernels from representative participants: win–stay/lose–shift, alternation, perseveration, and win–shift/lose–stay. Callout titles report the number of participants assigned to each pattern.

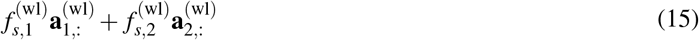

where 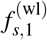 and 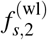 are the subject’s scores for reinforcement learning (RL) and choice bias, respectively, and 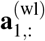, 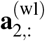 are the shared history components corresponding to the two dimensions. The scatter plot of participant scores (Fig. 8b) reveals that while most individuals exhibit a positive RL strategy (i.e., positive score 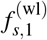 on the first history dimension), they are broadly distributed along the choice bias axis in participant-score space (i.e.,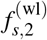.

Most people exhibit a Strong *Win →Stay* and *Loss →Shift* with little outcome–independent bias with a small 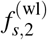 (Fig. 8c). Participants represented by the red dot in the top-`Right` quadrant are likely to be strong learners with a powerful alternation bias (Fig. 8d). For this group, the two influences align after a loss—the RL signal encourages them to switch, and their alternation bias also encourages them to switch, resulting in a very strong lose-shift tendency. After a win, however, the influences compete: the RL signal encourages them to stay, while their alternation bias pushes them to switch. Conversely, participant represented by the green dot in the bottom-`Right` quadrant are strong learners with a perseveration bias (a tendency to repeat choices). After a win, both the RL signal and their repetition bias encourage them to stay, creating a robust win-stay behavior. After a loss, these influences compete: the RL signal promotes a shift, but the bias promotes staying. This conflict can dampen or even override the lose-shift tendency, leading to behavior that is slow to adapt to negative feedback (Fig. 8e). Lastly, there are 6 people (Fig. 8f) shows a negative RL strategy (i.e., win-shift/lose-stay).

### 3.4 Replication with new data

We also validate our results with a newly collected, replication dataset from 102 participants. Following the same procedure as before, we ran the model across 121 training configurations, generated from 11 independent train/validation splits and 11 random weight initializations. The results replicated the same patterns identified in the original dataset. A total of six dimensions were discovered for the current data structure, including two dimensions for the temporal weighting kernel (See Fig. 9 I), two for the history kernel, and one dimension each for the side bias and random noise. First, the scaling factor did not vary much from before with a median of 0.347 (IQR = 0.178; *Q*_1_ = 0.244, *Q*_3_ = 0.421), indicating similar between-subject variability (See Fig. 9 II). Next, different from before, participants exhibited an overall numerically `Left` side bias but not significantly different from 0 *W* = 2347.00, *p* = .351, *Mdn* = *−* 0.244, *IQR* = 1.541, *N* = 102. Among all of them, 42 participants exhibited a `Right` side bias estimate and 60 exhibited a `Left` side bias estimate at the individual level (See Fig. 9 III). The temporal deviation is still captured by a bump-recency component and a primacy component, and individual temporal deviations can be clustered into 5 groups, including Bump, Primacy, Recency, Bookending and Flat (See Fig. 9 IV). Similarly, one reinforcement learning component (See Fig. 9 V) and one choice component was discovered under the history effect. Individuals can also be summarized by win-stay/lose-shift, alternation, perservation and win-shift/lose-stay strategy.

**Figure 9.**
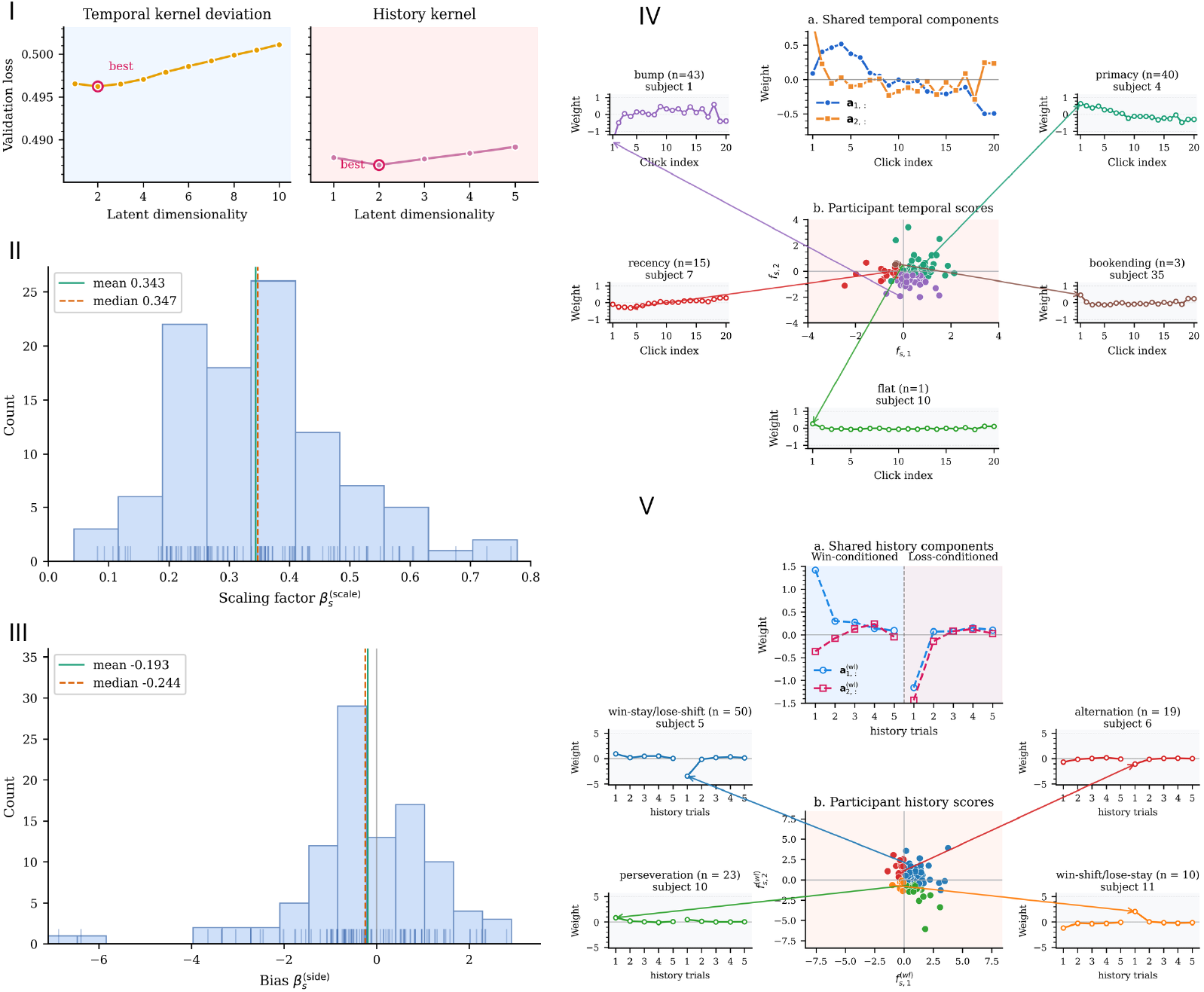
Results from validation dataset. I). Validation loss as a function of the number of latent dimensions for the temporal weighting kernel and the history kernel on the newly collected dataset. Each point shows the average validation loss over 121 training loops. Consistent with the original dataset, the minimum validation loss in both cases occurs at two latent dimensions, indicating a validation-selected dimensionality of *d*_*T*_ = *d*_*H*_ = 2 for temporal weighting and history dependence. II) and III). Distribution of estimated side bias and scaling factor for the newly collected dataset. Bars show the histogram of individual bias parameters, with rug marks indicating single participants. The mean/median bias are -0.193 and -0.243 respectively; The mean/median scaling are 0.343 and 0.347 respectively. IV). Shared temporal components, participant scores, and individual temporal kernel deviations in a two-dimensional representation for the newly collected dataset. (a) Per-click weights for the two shared temporal components. (b) Participants’ *f*_*s*,1_ and *f*_*s*,2_ scores; colors indicate clusters associated with distinct temporal weighting strategy. (b, callouts) Example temporal kernel deviations from individual participants representative of each cluster: bookending, primacy, recency, bump-shaped, and flat. The overall structure reproduces that observed in the original dataset, with the same qualitative temporal weighting kernels recovered. Panel titles report the number of participants assigned to each pattern. V). History latent space and example history kernels for the newly collected dataset. (a) Weights over the preceding five trials for the two shared history components. (b) Participants’ 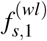 and 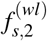 scores; colors indicate clusters associated with distinct history strategies. (b, callouts) Example kernels from representative participants: win–stay/lose–shift, alternation, perseveration, and win–shift/lose–stay. Consistent with the original dataset, the same qualitative history-strategy structure is observed.

## 4 Discussion

We investigated whether the diverse ways individuals depart from optimal behavior in perceptual decision-making can be explained by a small set of shared computational components. In the Bernoulli clicks task, out-of-sample prediction reveals a six-dimensional representation for individual differences: two dimensions of temporal weight-ing and two dimensions of history dependence, together with participant-specific side bias and signal-to-noise ratio. Similar structures emerged in an independent replication dataset. These findings suggest that apparently distinct decision strategies can be understood as different combinations of shared computational components. By estimating how participant-specific weightings of these components shape the decision variable on each trial, the model directly connects the macroscopic organization of behavioral differences across people to the trial-level computations that generate choices within each person.

Combinations of two shared temporal components captured primacy, recency, middle-weighted, bookending, and near-flat kernels. Within this representation, these kernels occupy different regions of a continuous two-dimensional space rather than falling into discrete strategy types. This continuity also explains how the same observer can shift between patterns, such as appearing primacy-like in one block and bump-shaped in another as experimental conditions change [23] or internal states drift [31, 35]. This picture complements existing accounts in which parameter variation within a common computational architecture—such as dynamic divisive normalization [31] or hierarchical approximate inference [16, 22]—produces diverse temporal weighting patterns. That is, our results add a population-level constraint to these accounts: a model of temporal integration should explain both the range of observed kernels and the low-dimensional organization of their variation across individuals.

The learned temporal space also offers a clear way to test specific mechanistic hypotheses. For example, leaky accumulation reduces the weight of earlier evidence, whereas early commitment to a decision reduces the weight of later evidence [17, 23]. Because different dynamical models can produce similar individual kernels, fitting single-participant kernels alone often cannot distinguish the underlying processes. Instead, we can ask whether varying the parameters of a candidate model reproduces the two-dimensional population space observed here. Experimental manipulations can then test whether changing evidence reliability, stimulus duration, or task demands moves participants through this space along the directions predicted by each model [23].

History dependence revealed a complementary two-dimensional history kernel space. Previous studies in the clicks paradigm have observed that both outcome-dependent (win–stay/lose–shift) and outcome-independent (repetition or alternation) history biases shape choices in rats [15, 21] and humans [18], a pattern corroborated across motion, orientation, and other sensory decision tasks [21, 34, 35]. Here we provide empirical quantitative evidence for this two-dimensional structure learned jointly across participants. The need for two dimensions shows that individual variation in history biases cannot be explained by a single heuristic whose overall strength varies across people; instead, participants systematically differ in how they balance outcome-dependent and outcome-independent influences.

Several mechanisms may underlie these two history components. A classic view understands history effects as driven by belief updates about environmental stationarity or non-stationarity [14, 15, 41]. Under perceived volatility, an optimal learner place greater weight on recent outcomes, strengthening the win–stay/lose–shift component. In contrast, persistent beliefs in environmental stationarity down-weight recent outcomes in favor of habitual staying or alternating). From an exploration–exploitation perspective, the win–stay/lose–shift component represents an adaptive policy in which win–stay implements exploitation and lose–shift samples an alternative, corresponding to undirected exploration [19, 37]. In this view, the outcome-independent choice bias component reflects a tendency to repeat previous choices, stabilizing exploitation runs, whereas reduced repetition bias or increased alternation may favor undirected exploration. Motor or effector hysteresis may also contribute to repetition bias, as prior hand use can bias current choices toward the same effector [36], especially when action costs are similar [30].

Beyond characterizing cognitive strategies, learning this structure directly from trial-by-trial choices confers a substantial methodological advantage: dramatic parameter efficiency and built-in statistical regularization [33]. Standard computational modeling, by fitting individual subjects separately, decouples choice estimation from dimensionality reduction (or regularization). In this task, a model with decision noise, side bias, temporal integration and history kernels requires 32 parameters per participants or 4960 to fit the population. By contrast, the best-fitting neural network model has only 990 parameters for the entire population equivalent to around 6 per person. This ensures that the learned representation captures genuine trait variance rather than idiosyncratic measurement noise.

Overall, our results demonstrate that the geometry of individual differences can directly constrain mechanistic models of perceptual decision-making, revealing that diverse integration strategies and history biases arise from distinct weightings over a small set of shared computational components. This framework opens several avenues for future investigation. Within sensory evidence accumulation, it provides a principled way to test whether these twodimensional temporal and history kernel spaces generalize across sensory modalities [25, 27], continuous-time environments, and varying speed–accuracy regimes [10]. Beyond perceptual tasks, the same population-constrained approach could guide computational discovery in domains characterized by substantial individual heterogeneity, such as value-based economic choice and multi-armed bandit reinforcement learning [7]. Finally, integrating low-rank behavioral representations with physiological recordings—such as EEG, pupillometry [12, 13], or intracranial recordings [39]—could determine whether these latent axes map onto distinct neural signatures of evidence accumulation [26, 29], urgency, or arousal. While bridging latent behavioral traits and neural dynamics requires explicit observation models that account for measurement noise, predicting both trial-by-trial decisions and neural state transitions [24] from a unified, low-dimensional basis offers a powerful paradigm for uncovering the shared architecture of human cognition.

## 5 Methods

### 5.1 Participants

We analyzed two datasets. A total of 186 undergraduate students from University of Arizona took part in the experiment for course credit in the first dataset.^17^ 31 subjects were excluded due to poor performance (accuracy lower than 60%), 155 subjects’ data were used in the current research. The experiment was approved by the Institutional Review Board at University of Arizona and all participants provided informed consent. For the second dataset, 124 participants were recruited through Prolific. We excluded 22 subjects due to poor performance (accuracy lower than 60%). The experiment was approved by the Institutional Review Board at Georgia Institute of Technology and all participants provided informed consent.

### 5.2 Bernoulli clicks task

Participants made a series of auditory perceptual decisions. After the fixation for 1 second, they listened 20 clicks over 1 second. Clicks are either on “`Right`” ear or “`Left`” ear. Participants decided which ear received the most clicks. Participants received feedback for 500 ms after the response.Partipants made their response using a standard keyboard, and played until they either gave 500 correct responses, or reached 50 minutes of experiment time. Unlike the Poisson clicks task,^5^ clicks in our task were delivered every 50 ms, with a fixed probability (*P* = 0.55) that a click occurred in the “correct” ear. The correct side was assigned with a fixed probability of 50%).

### 5.3 Descriptive behavioral analyses

Figure 2 summarizes choice behavior in the first dataset. The calculations for each panel are described below. As described before, clicks were coded as *c*_*i,t*_ = *−*1 for `Left` and *c*_*i,t*_ = +1 for right, where *i* denotes click position and *t* denotes trial. Total evidence for each trial was 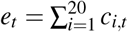, the number of `Right` clicks minus the number of `Left` clicks. Choices were coded as α_*t*_ = *−*1 for `Left` and α_*t*_ = +1 for right.

#### Panel A: Optimal-observer choice curve

The optimal curve follows the optimal decision rule in Eq. 2: choose `Right` when *e*_*t*_ *>* 0 and `Left` when *e*_*t*_ *<* 0. When *e*_*t*_ = 0, either response is equally likely. Thus, the probability of choosing `Right` is 0, 0.5, or 1 for negative, zero, or positive evidence, respectively.

#### Panel B: Human choice curve

For each participant, we grouped trials by total evidence and calculated the proportion of rightward choices at each observed evidence value from *−* 10 to +10 clicks. At each value, we then averaged these proportions across participants who had at least one trial with that evidence value. Each contributing participant received equal weight. The plotted points are these group means, and the lines connect adjacent points; no sigmoid was fitted.

#### Panel C: Model-free side bias

We used trials containing equal numbers of `Left` and `Right` clicks, to measure each participant’s empirical side preference. For participant *s*, we calculated the proportion of rightward choices on these trials and subtracted 0.5:

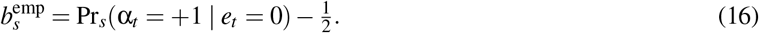

A positive value indicates a rightward preference, a negative value a leftward preference, and zero no preference. The histogram contains one bias value per participant with at least one zero-evidence trial. This empirical measure is distinct from the side-bias coefficient estimated by regression.

#### Panel D: Choice-triggered reverse kernel

The reverse kernel describes whether evidence at a given click position is more rightward on trials ending in a `Right` choice than on trials ending in a `Left` choice, after subtracting a task-derived reference. We calculated this descriptive measure at each of the 20 click positions in four steps.

First, derive the reference mean from the task probabilities. The baseline comes from how clicks were generated. For a trial with a `Right` target, each click was rightward with probability 0.55 and leftward with probability 0.45. Because `Right` and `Left` clicks were coded as +1 and *−* 1, respectively, the expected signed click was the probability-weighted average of these two values. The probabilities were reversed for a `Left` target:

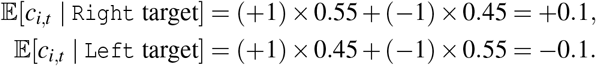

These expected values are the same at every click position because the generating probabilities were constant through-out the trial.

Second, express each click relative to the reference baseline. For the actual calculation, the reference sign was assigned from the majority side of the click sequence, sign(*e*_*t*_), rather than a separately recorded stimulus-generating target. We excluded equal-evidence trials (*e*_*t*_ = 0) because there was no majority side. Each remaining trial therefore had a reference of +0.1 for a `Right` majority or *−* 0.1 for a `Left` majority. We subtracted this reference from each click:

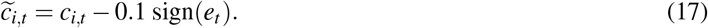

On a right-majority trial, a `Right` click gives 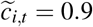 and a `Left` click gives 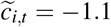. Thus, 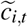 expresses each click as a signed deviation from the baseline.

Third, compare the deviations between the two choice groups. Within each participant, we divided the retained trials into right-choice and left-choice groups. At each click position *i*, we summed 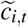 over right-choice trials and divided by the number of those trials. We repeated this calculation for left-choice trials, then subtracted the left-choice mean from the right-choice mean:

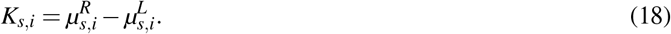

Here, 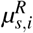 and 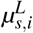 are participant *s*’s mean baseline-corrected clicks, 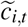, at position *i* on right-choice and left-choice trials, respectively. A positive *K*_*s,i*_ means that the deviation at position *i* was, on average, more rightward before `Right` choices than before `Left` choices; zero means that the two group means were equal. Comparing *K*_*s,i*_ across positions gives the temporal pattern of this choice-related difference.

Finally, average across participants and quantify uncertainty. We first obtained a complete 20-position kernel for each of the 155 participants, then averaged their kernels with equal participant weights:

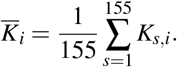

The plotted line shows 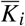. At each position, the shaded band is the pointwise normal-approximation 95% confidence interval, 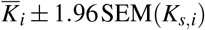, where the SEM is the sample standard deviation of the 155 participant-specific kernel values divided by 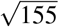. The interval therefore summarizes uncertainty in the across-participant mean at that position.

#### Panel E and F: Choice curves after a win

We sorted trials within each participant as they appeared in the task. A previous trial was classified as a win when the participant’s choice matched the correct answer. We selected current trials that followed a win and divided them according to whether the previous choice was `Left` or right. Within each group, we calculated each participant’s proportion of current rightward choices at each current trial total evidence value from -10 to +10. After that, we then averaged these proportions across participants. This produced the previous `Left` and previous `Right` curves in panel E. The first trial of each participant and trials immediately following an equal evidence trial were excluded. These same rules apply to panel F.

### 5.4 Statistical analyses for Figure 3

Statistical analyses used the parameters fitted separately to each of the 155 participants in Fig. 3. We analyzed the raw side-bias, temporal-deviation, and history coefficients before multiplication by the participant-specific scaling parameter. The top row of the figure shows the equally weighted mean across participants, with error bars of *±*1 SEM; the SEM was the sample standard deviation across participants divided by 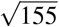. The scaling parameter and the two example participants were shown descriptively.

### Temporal weighting

To test whether temporal weighting varied across the trial, we entered each participant’s 20 click weights into a one-way repeated-measures ANOVA, with click position as the within-participant factor. The null hypothesis was that the population mean weight was the same at all 20 positions. The calculation used 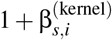; adding one to every value leaves this test unchanged. We applied the Greenhouse–Geisser correction to both degrees of freedom and calculated the *p* value from the corrected degrees of freedom. The effect size was partial 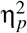, calculated as the click-position sum of squares divided by the sum of the click-position and residual sums of squares.

We also tested which positions were over- or underweighted relative to the baseline. At each position, a two-sided one-sample *t*-test compared the 155 temporal-deviation coefficients, 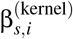, against zero. These 20 tests formed one Benjamini–Hochberg false-discovery-rate (FDR) correction family. Cohen’s *d*_*z*_ was calculated as the mean coefficient divided by its sample standard deviation across participants.

#### Side bias

Each participant contributed one side-bias coefficient,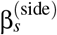. We used a one-sample Wilcoxon signed-rank test against zero, with the one-sided alternative that side-bias coefficients were shifted toward positive values, indicating a rightward bias.

#### History effect

We first averaged each participant’s five win-history coefficients across 5 previous trials separately, giving one win-history mean per participant. We did the same for the five loss-history coefficients. Separate one-sample Wilcoxon signed-rank tests compared these 155 participant means against zero. The win-history test used the one-sided alternative of a positive shift, corresponding to win–stay behavior; the loss-history test used the one-sided alternative of a negative shift, corresponding to lose–shift behavior. These tests assessed the average effect across the five history trials, rather than each trial separately.

#### Signed-rank statistics and significance threshold

For all three Wilcoxon tests, zero values were omitted, the absolute nonzero values were ranked, and ties received average ranks. The reported statistic *W* was the sum of ranks for positive values. Effect sizes were signed rank-biserial correlations, *r* = (*R*_+_ *− R*_*−*_)*/*(*R*_+_ + *R*_*−*_), where *R*_+_ and *R*_*−*_ are the positive and negative rank sums. Positive *r* indicates a predominance of positive coefficients and negative *r* a predominance of negative coefficients. We also reported the across-participant mean and median to describe the magnitude of each effect. Statistical significance was assessed at *p <* 0.05, using the Greenhouse–Geisser-corrected *p* value for the ANOVA and FDR-adjusted *p* values within the two multiple-testing families.

### 5.5 Model derivation

For each trial *t*, the click sequence is denoted by

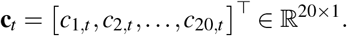

In the optimal model, all clicks are weighted equally. Therefore, the optimal accumulator of the sensory evidence for each subject is

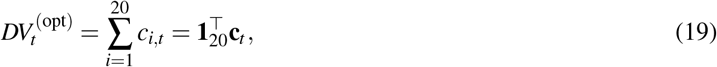

where

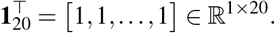

However, individual’s behavior are noisy, we introduce the scaling parameter 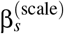 to express this noisy computation. In this case, subject does not have to weigh each click as 1. It is constraint to be positive and produced from subject identity:

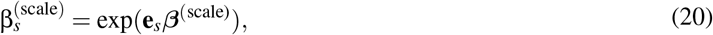

where

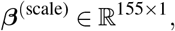

which maps the one-hot encoded subject vector

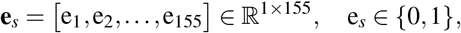

to a subject specific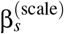.

In this optimal case, 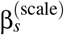 approaches infinity, producing deterministic choices from the sign of the accumulated evidence, because the 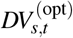 is mapped to choice with a logistic link:

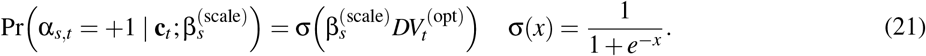

The first suboptimality we introduced was a subject-specific side bias. This term captures a general tendency of subject *s* to choose `Left` or `Right`, independent of the click evidence. The side bias is produced from the one-hot subject vector:

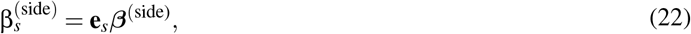

where

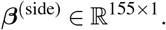

The next suboptimality we introduced was a subject-specific temporal integration deviation. In the optimal model, the baseline click weight at every temporal position is 1. For subject *s*, the temporal kernel deviation is written as

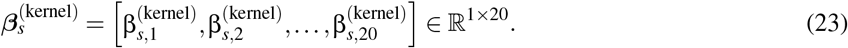

Each element 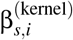 captures how subject *s*’s weight at click position *i* deviates from the optimal baseline weight (Fig. 10).

**Figure 10.**
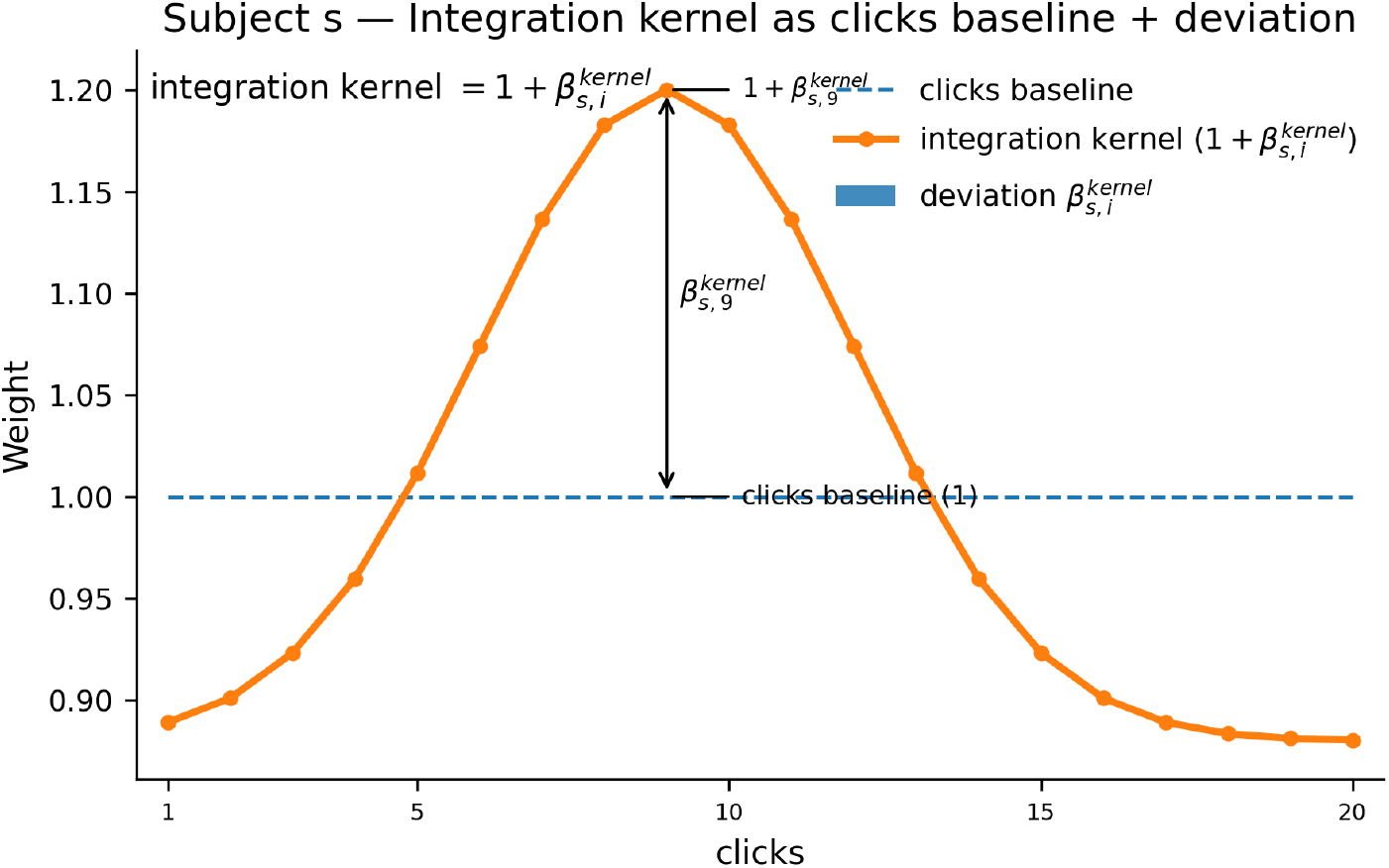
Example decomposition of the temporal integration kernel into a uniform baseline and a temporal kernel deviation. The x-axis indexes the 20 click positions in time, and the y-axis shows their decision weight. The dashed horizontal line denotes the clicks baseline, corresponding to perfectly uniform weighting of all clicks. The orange curve shows the subject’s integration kernel, given as 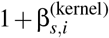. This bump-shaped curve indicates that this hypothesized subject overweights middle clicks and underweights early and late clicks. The vertical arrow at click 9 illustrates the deviation 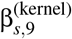 from baseline; analogous deviations at each position capture the full kernel shape.

Therefore, the full temporal integration weights for subject *s* are

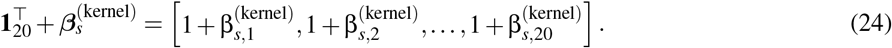

To estimate the latent dimensionality of individual temporal kernel deviations, we apply a low rank factorization to the kernel parameter:

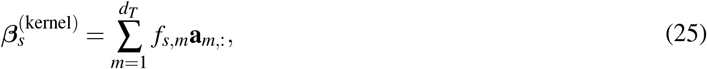

where **a**_*m*,:_ *∈ ℝ* ^1*×*20^ is the shared temporal component corresponding to latent dimension *m*, and *f*_*s,m*_ is subject *s*’s score on that dimension.

Coupling equation (25) with the click sequence gives the accumulated evidence on trial *t*:

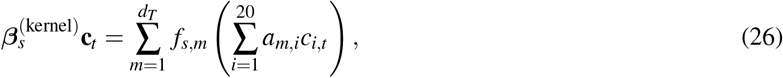

written in vector representation form

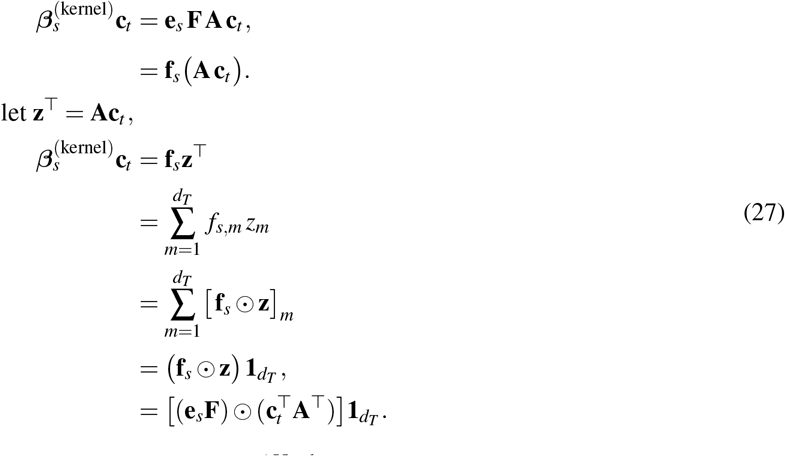

Here, **f**_*s*_ = **e**_*s*_ **F** is the output from the subject layer, 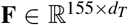 is the matrix of the subject layer, denoting participant scores, with (*s, m*)-th entry *f*_*s,m*_, and **e**_*s*_ *∈* ℝ^1*×*155^ is the one-hot vector selecting subject *s*. The matrix 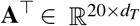 contains the temporal components: its *m*-th column is 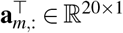, equivalently (**A**^*⊤*^)_*i,m*_ = *a*_*m,i*_. Consequently, **e**_*s*_**F** and 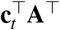 are both row vectors in 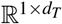. The operator *⊙* denotes elementwise multiplication, and 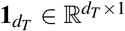 sums the resulting contributions across the *d*_*T*_ temporal components.

Adding this temporal deviation and bias term gives the following:

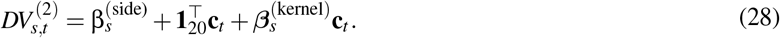

Substitute equation (27) into equation (28), we obtain the vector form of the model:

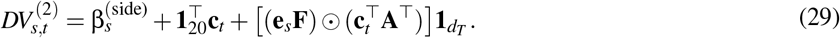

And this is our first model, used to estimate the latent dimensionality of temporal kernel deviations. In a fully unconstrained model, the temporal kernel deviation coefficients for all *S* = 155 subjects for 20 clicks would form an unconstrained coefficient matrix ***β***^(kernel)^ *∈ ℝ* ^155*×*20^. Our model essentially constrains these weights to a low-dimensional temporal kernel space by enforcing ***β***^(kernel)^ = **FA**, where 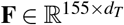 contains the participant scores and 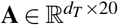 contains the shared temporal components. The temporal latent dimensionality *d*_*T*_ is selected by comparing validation loss across models with different numbers of shared temporal components.

The training procedure follows an iterative approach to select the temporal latent dimensionality. Each latent dimension is represented by a shared temporal component and its participant scores, stored as corresponding columns of **A**^*⊤*^ and **F**, respectively. In the neural-network implementation, each dimension corresponds to one unit in each of the click and subject layers. The process begins with an initial neuron count of *d*_*T*_ = 1 and a placeholder for the best loss set to an arbitrarily high value (Algorithm 1), after each iteration, the number of neurons in both the click and subject layer increased by one. The latest model inherits the weight from the previous well-trained model and only optimizes the new neuron’s weight during training. In each iteration, the model is trained and evaluated on a validation set to compute its corresponding loss, *L*. If *L* is lower than the current best-recorded loss minus the improvement threshold, the best neuron count and loss are updated accordingly. Otherwise, the previous best configuration remains unchanged. This process continues until the number of neurons reaches *d*_*T*,max_ = 10. This iterative approach ensures that every dimension fit to the residual noise, and each new dimension is approximately fitted to the residual noise that is not well explained by previous dimensions, and all configurations up to the predefined limit are explored, providing a systematic evaluation of model performance.

Finally, we incorporated trial-history effects using win–stay and lose–shift terms from the previous five trials. Let α_*t−j*_ *∈ {−*1, 1*}* denote the previous choice or behavior on trial *t − j*, where *−*1 indicates `Left` and 1 indicates `Right`. Let *w*_*t−j*_ indicate whether trial *t − j* was correct, and let *l*_*t−j*_ indicate whether trial *t − j* was incorrect. We write

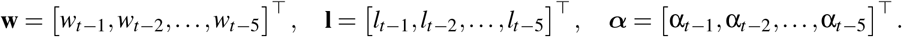

The unfactorized precvious–win contribution is

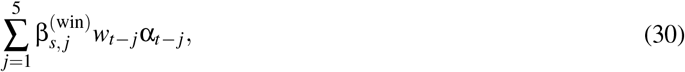

and the unfactorized previous–lose contribution is

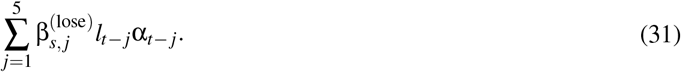

As with the temporal coefficient matrix, we represent the joint history coefficient matrix using a low-rank factorization. For the win-conditioned choice-history term, we write

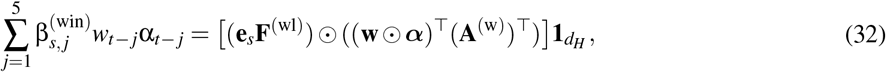

where

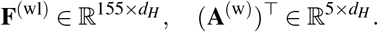

Similarly, for the loss-conditioned choice-history term, we write

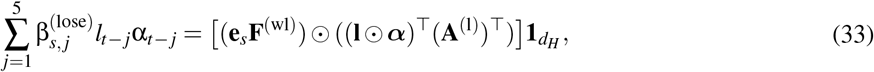

where

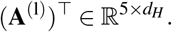

The final decision variable is:

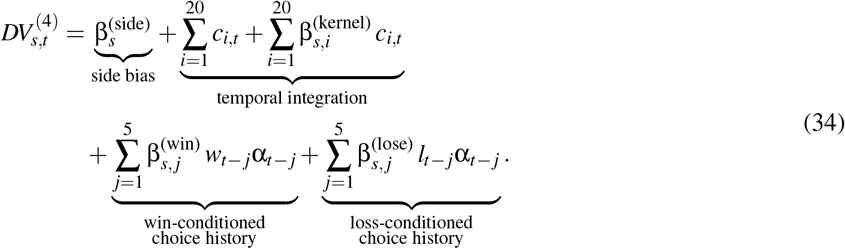

Matching our implementation of the neural network, we have:

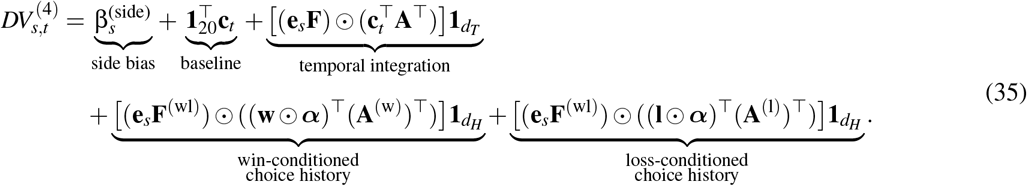

The final history model uses a shared subject-score matrix **F**^(wl)^ for the win and loss terms. Writing **A**^(wl)^ = [**A**^(w)^ **A**^(l)^], the concatenated subject-by-history coefficient matrix is

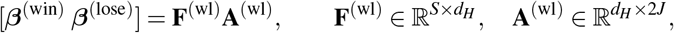

where *d*_*H*_ is the joint history dimensionality and *J* = 5. Thus, the concatenated matrix has rank at most *d*_*H*_, and each subject score jointly scales the win and loss portions of a history component. Algorithm 2 adds these joint components after fitting the temporal model. Each expansion adds one shared subject-score column and one history-basis row containing both the win and loss weights.

### 5.6 Training algorithm

### 5.7 Objective function and Generalized Low-Rank Regularized Model

Motivated by the generalized low-rank modeling viewpoint—where Principal Component Analysis (PCA) can be framed as a low-rank approximation under squared loss, and this principle extends by pairing low-rank structure with a loss matched to the observation model (e.g., logistic loss for binary data)^33^—we adapt this idea to behavioral choice modeling by imposing a low-rank structure on the subject × feature coefficient matrix **W**. Our work models the behavioral choice data as a low-rank factorized logistic regression, with the goal of predicting subjects’ binary choices while condensing their decision strategies (and suboptimalities) into a shared, low-dimensional space.

Let α_*s,t*_ *∈ {−*1, +1*}* represent the choice made by subject *s* on trial *t*, with +1 denoting a rightward choice and *−*1 a leftward choice. As described above in section 5.5, we model the choice probability of choosing `Right`, 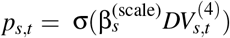, as a logistic regression model (see equation 34). Thus the parameters **F, A, *β***^(scale)^, ***β***^(side)^, **F**^(wl)^, **A**^(wl)^ are recovered from the behavioral data by solving

#### Algorithm 1 Iterative Expansion to Select Temporal Latent Dimensionality *d*_*T*_

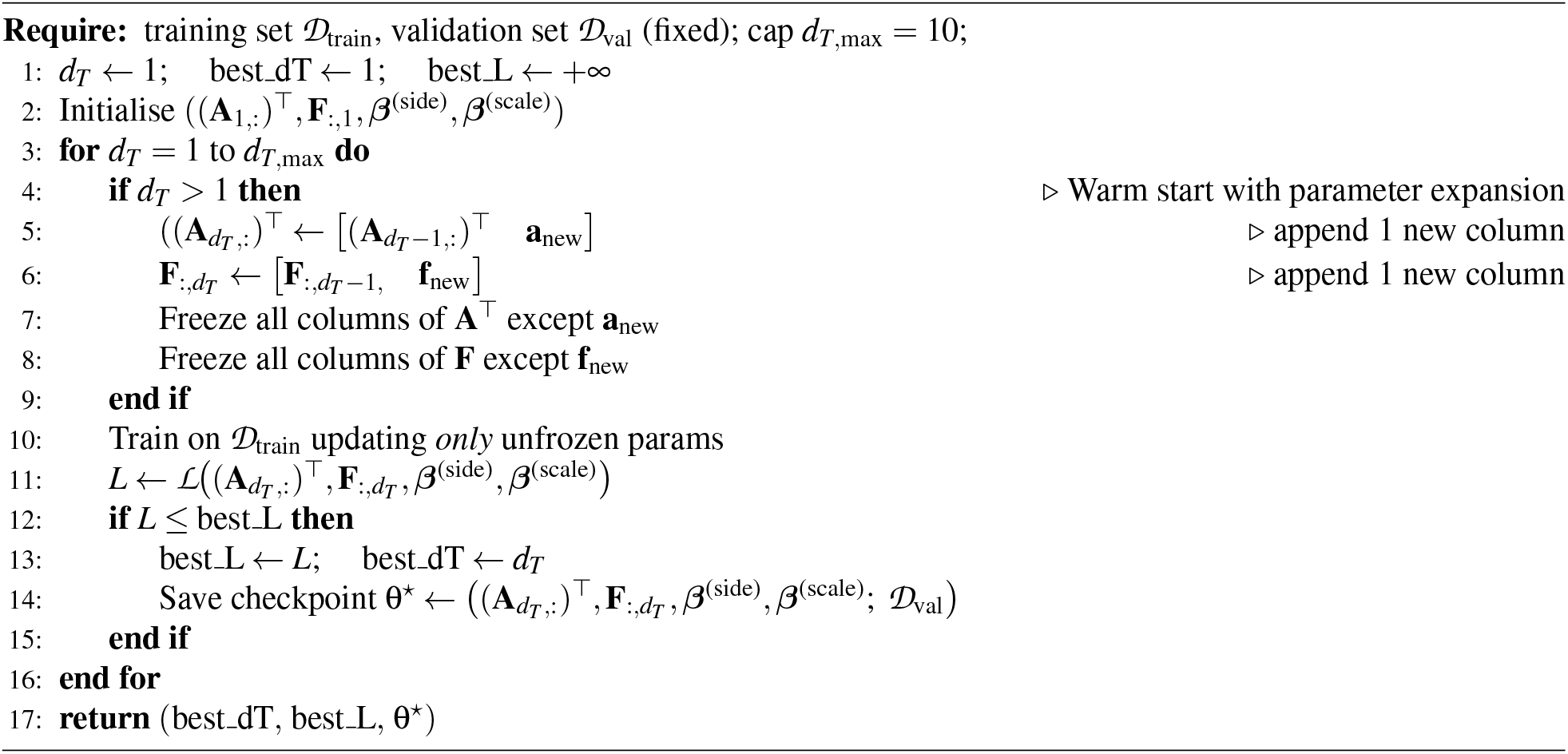

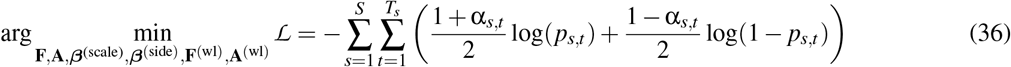

where *S* is the number of subjects and *T*_*s*_ is the number of trials for subject *s*.

To prevent noise inflation typical of independent per-subject modeling, we constrain the temporal and joint history coefficient matrices to have rank at most *d*_*T*_ and *d*_*H*_, respectively. This structural regularization factorized the integration and history kernels into shared bases and individual scores, resulting in a significant decrease in the effective degrees of freedom.

The robustness of the inferred latent representation arises from the coupling of the gradients of the above objective function across the population of subjects. To illustrate, 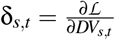 denote the scalar error signal for a specific trial. Then the update rule for the gradient descent for the shared temporal basis matrix **A**_*click*_ aggregates error signals via the outer product of the participant scores 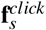 and the stimulus **c**_*t*_ :

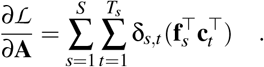

The key is that the gradient summation 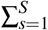 is over the population of subjects, so subject specific measurement noise from individual subjects is averaged out, and ensures that the updated learning of **A**_*click*_ incorporates shared, structural population-level signals. Then, by iteratively expanding the latent space (*d*_*T*_ from 1 to 10 and *d*_*H*_ from 1 to 5) and selecting the value of *d*_*T*_ /*d*_*H*_ that minimizes held-out validation loss, the ranks of the temporal and joint history coefficient matrices are bounded by *d*_*T*_ and *d*_*H*_, respectively. This cross-validated rank selection ensures that the learned representation captures stable, generalizable trait variance rather than measurement noise.

### 5.8 Hyperparameters

The model is trained using the following hyperparameters. Training is performed with a batch size of 32 and a fixed learning rate of 0.01 over a maximum of 5,000 epochs. In addition, 15% of the data is held out for validation and another 15% of the data is for evaluation (i.e., test set). Early stopping is implemented with a patience of 150 epochs for temporal deviation and 100 epochs for history kernel. In addition, an improvement threshold of 0 was set, with the waiting counter initialized to 0. `BCEWithLogitsLoss` is used to compute the loss. The training process consists of 11 different data splits and 11 different random initializations of weight parameters. This results in a total of 121 unique training loops. All code and data can be found at https://osf.io/2vcsx/.

#### Algorithm 2 Select the Rank of the Joint Win–Loss History Coefficient Matrix

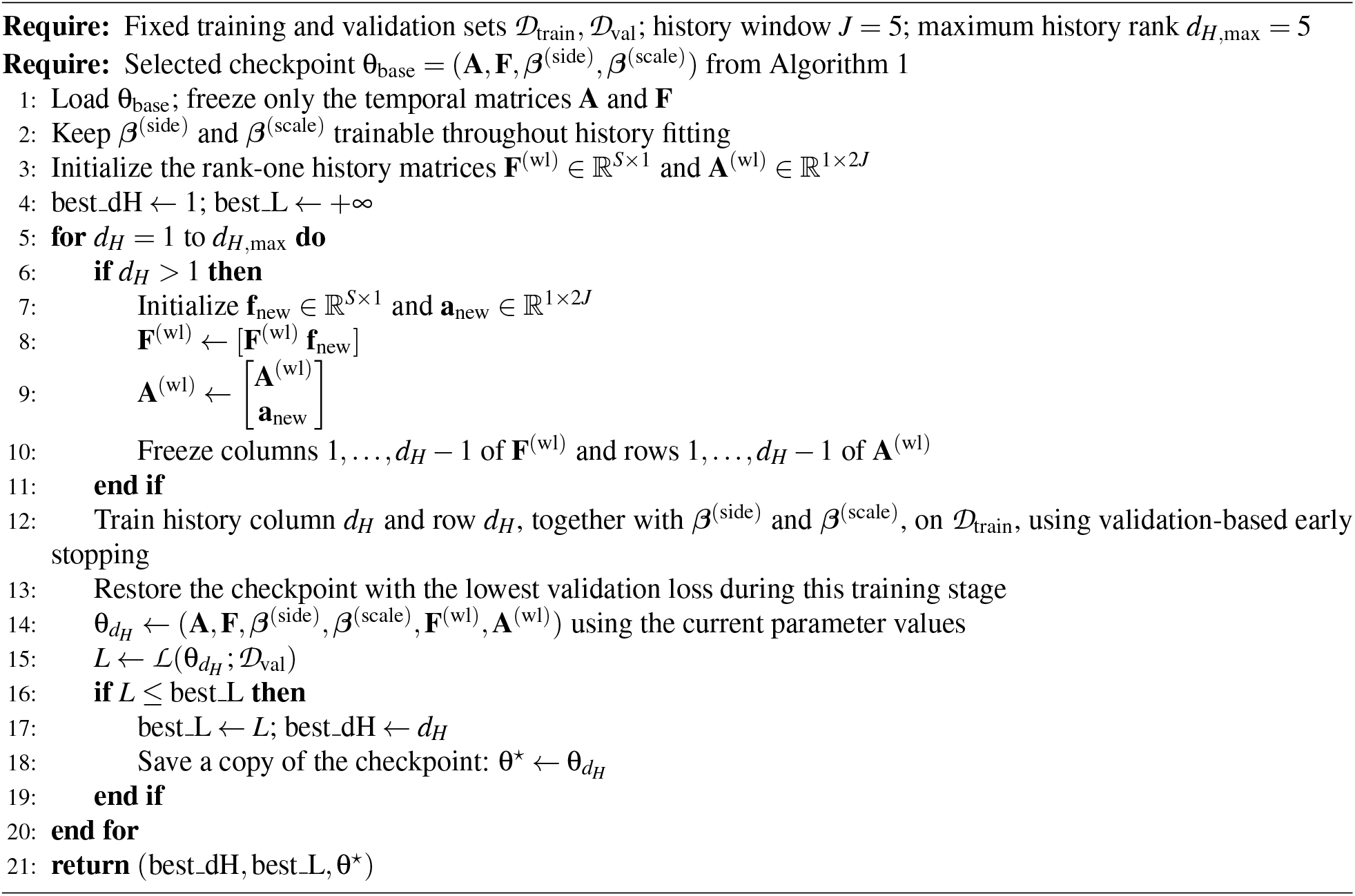

**Table 1:** Notation used in the Bernoulli Clicks Task models.

| Notation | Type | Definition |
|---|---|---|
| *s* | Subject index | Index identifying an individual participant. All parameters with subscript *s* are estimated separately for each subject. |
| *t* | Trial index | Index identifying the current trial. |
| *i* | Click-position index | Index for the temporal position of a click within a trial, with *i* = 1, …, 20. Each trial contains one click in each of twenty temporal bins. |
| *j* | History-lag index | Index for how many trials in the past an outcome occurred, with *j* = 1, …, 5. For example, *j* = 1 denotes the immediately preceding trial. |
| **c**<sub>*t*</sub> | Click vector | Vector containing the full sequence of clicks presented on trial *t*: **c**<sub>*t*</sub> = [*c*<sub>1,*t*</sub>, *c*<sub>2,*t*</sub>, …, *c*<sub>20,*t*</sub>]. |
| *c*<sub>*i*,*t*</sub> | Signed click | The click presented at temporal position *i* on trial *t*. It is coded as *c*<sub>*i*,*t*</sub> = +1 for a right-ear click and *c*<sub>*i*,*t*</sub> = −1 for a left-ear click. |
| $\sum_{i=1}^{20} c_{i,t}$ | Baseline evidence | Net right-minus-Left evidence on trial *t*. This is the decision variable used by the optimal observer before adding suboptimalities. |
| α<sub>*t*</sub> | Choice variable | Participant’s choice on trial *t*. For consistency with the history terms, it should be coded as α<sub>*t*</sub> = +1 for a Right choice and α<sub>*t*</sub> = −1 for a Left choice. |
| Pr(α<sub>*t*</sub> = right) or<br>*p*(α<sub>*t*</sub> = right) | Choice probability | Model-predicted probability that the participant chooses the Right side on trial *t*. |
| β<sub>opt</sub><sup>(scale)</sup> | Optimal inverse temperature | Inverse-temperature parameter of the optimal observer. In the deterministic optimal-observer limit, β<sub>opt</sub><sup>(scale)</sup> → +∞. |
| β<sub>*s*</sub><sup>(scale)</sup> | Evidence-scaling parameter | Subject-specific inverse-temperature parameter controlling sensitivity to the decision variable. Larger positive values correspond to more deterministic choices and greater sensitivity to accumulated evidence. |
| β<sub>*s*</sub><sup>(side)</sup> | Side-bias parameter | Subject-specific intercept added to the decision variable. Positive values indicate a bias toward Right choices, whereas negative values indicate a bias toward Left choices. |
| β<sub>*s*,*i*</sub><sup>(kernel)</sup> | Temporal kernel deviation parameter | Subject-specific deviation in the weight assigned to click position *i*. Because the baseline term already contributes *c*<sub>*i*,*t*</sub>, the effective weight on click *i* is 1 + β<sub>*s*,*i*</sub><sup>(kernel)</sup>. Positive values indicate overweighting and negative values indicate underweighting relative to uniform optimal weighting. |
| $\sum_{i=1}^{20} \beta_{s,i}^{(\text{kernel})} c_{i,t}$ | Temporal-deviation contribution | Contribution to the decision variable caused by nonuniform weighting of clicks across the twenty temporal positions. |
| *w*<sub>*t*−*j*</sub> | Win indicator | Binary variable indicating whether the choice on trial *t* − *j* was correct. It equals 1 for a win and 0 otherwise. |
| *l*<sub>*t*−*j*</sub> | Loss indicator | Binary variable indicating whether the choice on trial *t* − *j* was incorrect. It equals 1 for a loss and 0 otherwise. Under the present coding, *w*<sub>*t*−*j*</sub> + *l*<sub>*t*−*j*</sub> = 1. |
| β<sub>*s*,*j*</sub><sup>(win)</sup> | Win-history parameter | Subject-specific coefficient describing how a correct choice *j* trials earlier influences the current decision. A positive value promotes repeating the prior choice after a win, corresponding to win–stay behavior. |
| β<sub>*s*,*j*</sub><sup>(lose)</sup> | Loss-history parameter | Subject-specific coefficient describing how an incorrect choice *j* trials earlier influences the current decision. A negative value promotes switching away from the prior choice after a loss, corresponding to lose–shift behavior. |

## Acknowledgments

SFF acknowledges support from SAFIR Institute RP 3760.

